# Membrane-Free Alveolus-on-a-Chip via Biodegradable Scaffold Recapitulates Interstitial Mechanics, Immune Trafficking, and Aerosolized mRNA Delivery

**DOI:** 10.64898/2026.04.17.719302

**Authors:** Jae-Won Choi, Huy Hoang Nguyen, Abbas Jalili, Matthew Andersen, Si-Yang Zheng

## Abstract

The pulmonary alveolus is a highly specialized microenvironment where epithelial, interstitial, and immune components interact to maintain gas exchange and tissue homeostasis. *In vivo*, the air–blood barrier consists of an epithelial layer and a capillary endothelium separated by an ultrathin interstitium composed of extracellular matrix (ECM) and lung fibroblasts. However, most existing lung-on-a-chip platforms rely on permanent synthetic membranes, which fail to recapitulate the dynamic biological and mechanical properties of the native interstitium. Here, we present a membrane-free human alveoli-on-a-chip enabled by a biodegradable poly(lactic-co-glycolic acid) (PLGA) scaffold that is progressively replaced by fibroblast-derived ECM. This process reconstructs a biologically formed interstitial layer while preserving an alveolus-like dome architecture. The resulting system supports multicellular organization under air–liquid interface conditions, enabling epithelial barrier formation and surfactant-related phenotypes. Additionally, direct epithelial–fibroblast interactions enhanced surfactant-related phenotypes, as evidenced by increased SPC and LAMP3 expression. Importantly, we demonstrate that conventional rigid substrates promote fibroblast-to-myofibroblast differentiation, leading to elevated reactive oxygen species (ROS) production, increased epithelial cell death, and compromised barrier integrity. In contrast, the membrane-free PLGA system mitigates stiffness-driven myofibroblast activation, preserving epithelial viability and maintaining barrier function. These findings highlight the critical role of interstitial mechanics in regulating alveolar homeostasis and reveal limitations of conventional membrane-based platforms. The platform further enables chemokine-driven monocyte migration across the alveolar barrier, recapitulating key immune trafficking processes observed *in vivo*. In addition, aerosolized metal–organic framework (MOF) nanoparticles efficiently mediated mRNA delivery to epithelial and interstitial cells with minimal cytotoxicity and modest inflammatory responses. Together, this membrane-free alveoli-on-a-chip reconstructs essential structural, mechanical, and functional features of the human alveolar microenvironment and provides a physiologically relevant platform for studying pulmonary biology, fibrosis-related mechanisms, immune cell trafficking, and inhaled nanomedicine delivery.

## Introduction

The pulmonary alveolus is the fundamental structural and functional unit of the lung responsible for gas exchange (*1*). *In vivo*, the air–blood barrier consists of an alveolar epithelial layer and a capillary endothelial layer separated by an extremely thin interstitium that includes a continuous basement membrane with a thickness of less than 1–2 μm (*2, 3*). This specialized architecture minimizes diffusion distance and enables efficient gas exchange between inhaled air and circulating blood (*4, 5*).

Beyond serving as a structural spacer between epithelial and endothelial layers, the alveolar interstitium functions as a dynamic microenvironment that regulates tissue mechanics, cell–cell signaling, and immune responses (*6, 7*). The interstitial compartment contains lung fibroblasts, extracellular matrix (ECM) components, and interstitial fluid, forming a connective tissue layer that supports alveolar structure and function. Fibroblasts within this compartment exhibit an elongated morphology with cellular processes that extend through the surrounding matrix, forming interconnected networks that communicate with epithelial cells, endothelial cells, and neighboring fibroblasts through gap junction–mediated signaling (*8*).

Structurally, the alveolar interstitium can be divided into two distinct regions. The thin portion is specialized for gas exchange, where the basement membranes of alveolar epithelial and capillary endothelial cells are fused, resulting in minimal diffusion distance and little connective tissue (*9*). In contrast, the thick portion contains a defined interstitial space populated by fibroblasts and ECM fibers such as collagen and elastin (*4, 10*). This region provides mechanical support to the alveolar wall and acts as a buffer for interstitial fluid and cellular interactions.

Within this interstitial niche, fibroblasts play essential roles in maintaining lung homeostasis. These cells synthesize key ECM components, including collagen I, collagen III, elastin, fibronectin, and proteoglycans, which collectively maintain the mechanical stability and elastic recoil of the alveolar wall during respiratory cycles (*11, 12*). In addition, fibroblasts regulate epithelial cell behavior through paracrine signaling, contributing to epithelial differentiation, barrier maintenance, and surfactant homeostasis. Following tissue injury, fibroblasts also participate in wound healing and ECM remodeling processes that restore tissue architecture (*13*). However, dysregulation of these fibroblast-mediated ECM remodeling processes can lead to pathological thickening of the interstitium, resulting in severe and progressive fibrotic disorders such as interstitial lung diseases (ILDs) and idiopathic pulmonary fibrosis (IPF) (*14*).

The alveolar interstitium further serves as a critical pathway for immune cell trafficking (*15*). During pulmonary inflammation or infection, circulating neutrophils and monocytes migrate across the vascular endothelium into the interstitial compartment and subsequently traverse the epithelial barrier to reach the alveolar airspace (*16*). This process is guided by chemokine gradients and coordinated interactions among endothelial cells, fibroblasts, and epithelial cells. Within the alveolar lumen, alveolar macrophages function as sentinel immune cells that recognize and phagocytose invading pathogens while orchestrating broader immune responses (*17–19*). In addition to these luminal macrophages, the alveolar interstitium harbors a distinct population of interstitial macrophages that are increasingly recognized for their vital roles in lung homeostasis and localized immune regulation (*20*).

Despite the importance of this complex multicellular microenvironment, current lung-on-chip platforms rely on permanent synthetic membranes that replace the biological interstitium with an inert artificial barrier (*21–27*). While these membranes provide mechanical support and permeability, they fail to recapitulate the dynamic biological properties of the native interstitium, particularly the fibroblast-derived ECM network and the associated cellular interactions that regulate tissue function and immune responses.

To address this limitation, we developed a membrane-free lung alveoli-on-a-chip that reconstructs key structural and functional features of the human alveolar microenvironment. Alveolar units form a largely polyhedral network with characteristic dimensions of approximately 200 μm. To emulate this physiological architecture, we fabricated biodegradable porous poly(lactic-co-glycolic acid) (PLGA) membranes that initially serve as a structural scaffold but gradually degrade during culture. As the scaffold degrades, it is progressively replaced by fibroblast-derived ECM, resulting in a biologically formed interstitial layer that more closely resembles the native alveolar tissue structure.

The reconstructed interstitial microenvironment also provides a unique platform for evaluating inhaled nanomedicine delivery under physiologically relevant conditions (*28*). In particular, metal–organic framework (MOF) nanoparticles have recently emerged as promising carriers for nucleic acid delivery due to their high cargo loading capacity and structural stability during aerosolization (*29, 30*). However, the biological responses and delivery efficiency of aerosolized nanocarriers remain difficult to evaluate using conventional *in vitro* systems (*31, 32*).

Here, we demonstrate that the membrane-free alveoli-on-a-chip supports multicellular alveolar tissue formation, enables chemokine-driven immune cell migration, and allows efficient aerosolized MOF nanoparticle–mediated gene delivery.

## Results

### Design of a biodegradable alveoli-mimetic membrane

To mimic the microenvironmental architecture of the pulmonary alveolus (Fig. 1A), we selected poly(lactic-co-glycolic acid) (PLGA) as the membrane material owing to its biodegradable and biocompatible properties (*33*), which enable the fabrication of physiologically relevant tissue models. Using this material, we fabricated alveoli-mimetic PLGA membranes with a highly porous architecture (Fig. 1, B and C). The engineered membrane exhibits an alveolar dome-like geometry with an interconnected porous network that facilitates molecular transport and cell interactions. Importantly, the membrane is designed to gradually degrade during culture, allowing it to be progressively replaced by ECM deposited by lung fibroblasts. As degradation proceeds, the fibroblast-derived ECM forms an interstitial layer and maintains a dome-shaped architecture with a diameter of approximately 200 µm, closely resembling the size and structure of native pulmonary alveoli.

**Figure 1.**
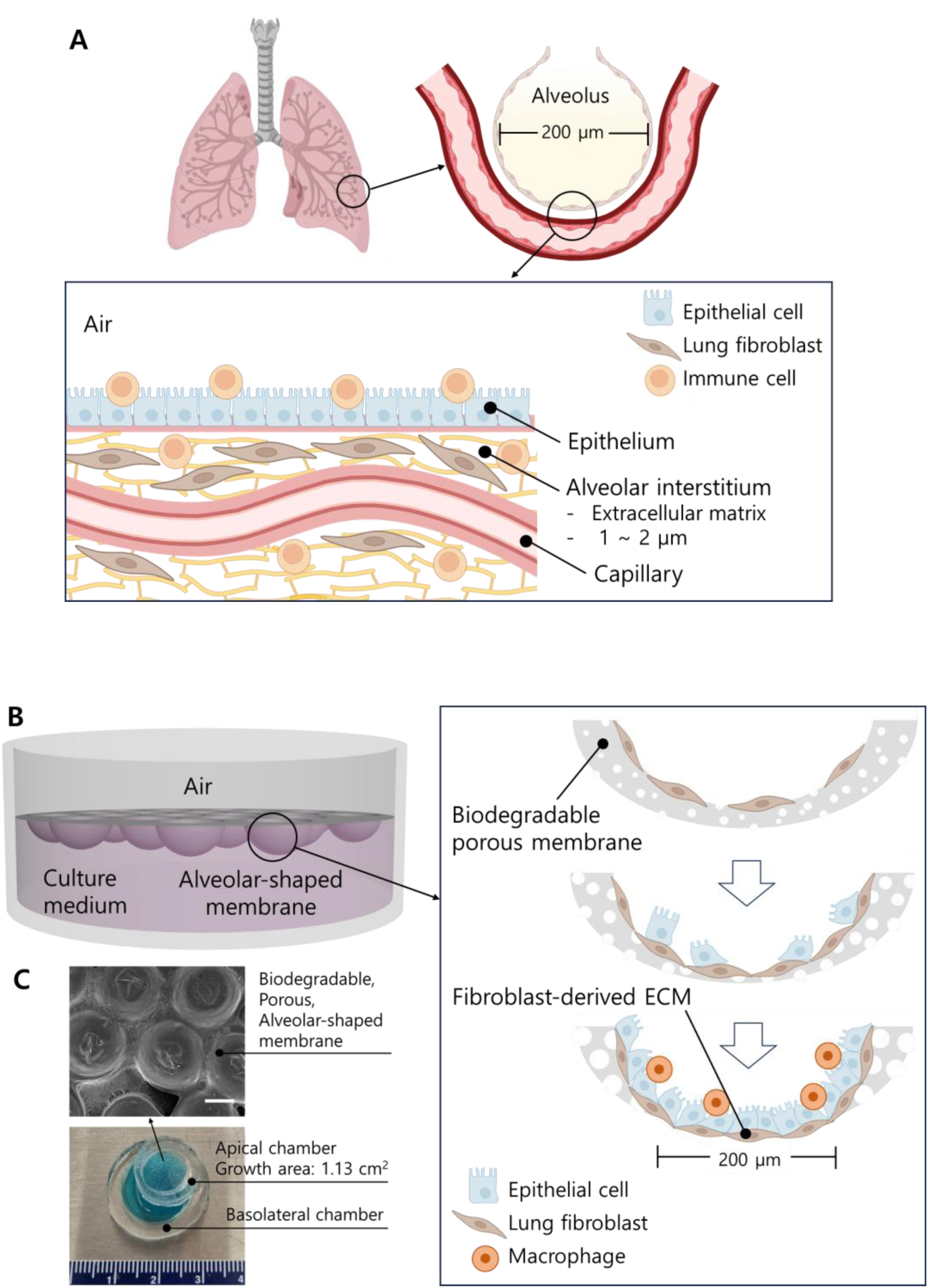
Biomimetic design of the membrane-free alveoli-on-a-chip. **(A)** Schematic illustration of the alveolar microenvironment. The pulmonary alveolus has an average diameter of approximately 200 μm and consists of an epithelial layer lining the air interface and an interstitial compartment composed of extracellular matrix (ECM) populated with lung fibroblasts. **(B)** Conceptual design of a membrane-free alveolus-on-a-chip platform based on a biodegradable membrane. The curved biodegradable membrane enables reconstruction of an alveolus-like architecture and supports the formation of a biomimetic alveolar microenvironment. **(C)** Photograph of the assembled chip showing the porous membrane integrated between the apical and basolateral chambers. SEM image showing dome-shaped alveolar microstructures formed on the porous biodegradable membrane. Scale bar: 200 μm.

On top of this interstitial-like layer, alveolar epithelial cells are cultured and subsequently exposed to air–liquid interface (ALI) conditions, enabling epithelial maturation and surfactant secretion, thereby more closely mimicking the physiological environment of the alveolus. Furthermore, the model enables the incorporation of immune cells, which can migrate across the epithelial and interstitial layers, allowing investigation of immune cell transmigration and multicellular interactions within a biomimetic alveolar microenvironment.

### Degradation-driven structural evolution of alveoli-mimetic PLGA membranes

An interstitium-mimicking membrane was developed using a dual-templated nonsolvent-induced phase separation (NIPS) strategy to fabricate poly(ε-caprolactone) (PCL) membranes (self-cited). Building on this approach, we extended the method not only to PCL but also to PLGA and employed PDMS molds containing curved microstructure arrays to engineer alveoli-mimetic membranes (Fig. S1), yielding a dome-like architecture that recapitulates the geometry of native alveoli (Fig. 2A and fig. S2A). Under aqueous physiological conditions, the alveoli-mimetic PLGA membranes underwent gradual degradation over a period of 10 days. During this process, the relatively thin central region of the alveolar dome preferentially degraded, resulting in the formation of a central pore. By day 7 of degradation, the diameter of this central pore reached approximately 200 μm, closely matching the average diameter of human alveoli (Fig. 2B) (*1*). A microporous architecture was formed on both PCL and PLGA membrane surfaces via NIPS, using camphene’s dendritic crystallization as a sacrificial porogen (Fig. 2C and fig. S2B) (*34*). After spin-coating the PLGA (or PCL) and camphene solution onto the mold, immersion in water, which acts as a nonsolvent for both PLGA (or PCL) and camphene, induced rapid solvent exchange. As acetone diffused outward and water penetrated into the solution, PLGA (or PCL) precipitated while camphene formed a dendritic crystalline network within the solidifying polymer matrix (fig. S3) (*35*). Subsequent freeze-drying sublimated the camphene, leaving behind voids that transformed the dendritic crystals into a porous structure defined by both phase separation and porogen templating. The initial surface porosity of the PLGA membrane was 19.5%, which is approximately 2.1-fold higher than that of conventional Transwell® membranes (Fig. 2D and fig. S4). As degradation progressed, the porosity of the PLGA membrane increased continuously, reaching approximately 43.5% by day 10. Concurrently, the average pore size increased from ∼1.12 μm to ∼3.98 μm (Fig. 2E). Mass-loss analysis further confirmed the progressive thinning and increasing surface porosity of the PLGA membranes during degradation, whereas the PCL membranes exhibited negligible degradation (fig. S2C). After 10 days in PBS, 3.71% of the initial mass had been lost, corresponding to an average degradation rate of approximately 3.7% per day, consistent with hydrolytic degradation (Fig. 2F).

**Figure 2.**
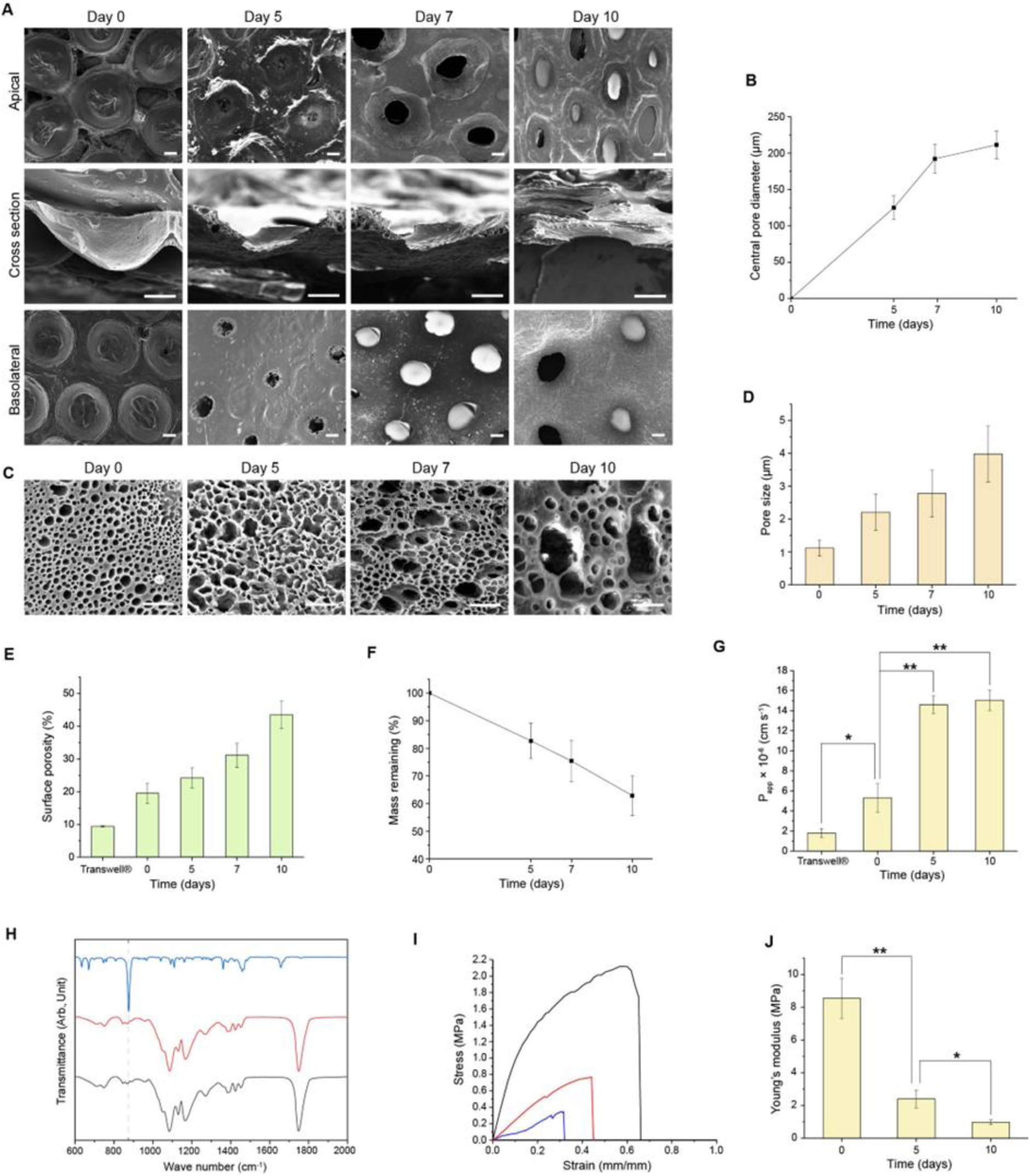
Degradation-driven structural, mechanical, and functional evolution of alveoli-mimetic PLGA membranes. **(A)** Representative SEM images of alveoli-mimetic PLGA membranes after degradation in PBS at 37 °C for 0, 5, 7, and 10 days. Scale bars: 100 μm. **(B)** Time-dependent changes in the diameter of central pores formed at the center of alveoli-mimetic PLGA membranes during degradation in PBS at 37 °C. (**C)** Representative SEM images showing the microporous structure of alveoli-mimetic PLGA membranes after degradation in PBS at 37 °C for 0, 5, 7, and 10 days. Scale bars: 10 μm. **(D)** Pore size of alveoli-mimetic PLGA membranes after degradation in PBS at 37 °C. **(E)** Surface porosity of Transwell® membranes and alveoli-mimetic PLGA membranes as a function of degradation time in PBS at 37 °C (n = 4). **(F)** Time-dependent mass loss of alveoli-mimetic PLGA membranes during incubation in PBS at 37 °C (n = 4). **(G)** Apparent permeability (P_app_) of 4 kDa dextran across Transwell® membranes and alveoli-mimetic PLGA membranes after degradation in PBS at 37 °C for the indicated time points (n = 4). **(H)** FT-IR spectra of camphene (blue), pure PLGA (red) and alveoli-mimetic PLGA membranes (black). **(I)** Representative stress–strain curves of alveoli-mimetic PLGA membranes after degradation in PBS at 37 °C for 0 (black), 5 (red), and 10 days (blue). **(J)** Young’s modulus of alveoli-mimetic PLGA membranes after degradation in PBS at 37 °C for 0, 5, and 10 days (n = 4).

### Chemical and mechanical properties of the PLGA membrane

PLGA membrane permeability, a key determinant of intercellular crosstalk and the transport of nutrients, growth factors, and signaling molecules, was markedly higher than that of the Transwell® membrane. The apparent permeability of the PLGA membrane was 3.0-fold greater prior to degradation and increased to 8.4-fold after 10 days of degradation compared with the Transwell® membrane (Fig. 2G). FT-IR spectra of the porous PLGA membrane showed a characteristic carbonyl stretching peak at 1750 cm⁻¹. In contrast, the characteristic camphene peaks corresponding to the C=C group at 1658 cm⁻¹ and aromatic C–H at 876 cm⁻¹ were absent, confirming the complete removal of the porogen (Fig. 2H) (*36*). The mechanical stability of the PLGA membrane during degradation was assessed by determining the apparent tensile modulus from the stress–strain curves (Fig. 2I). During hydrolytic degradation over 10 days, the modulus of the PLGA membrane progressively decreased from 8.55 MPa to 0.98 MPa (Fig. 2I). The decrease in tensile modulus was accompanied by reductions in ultimate tensile strength (UTS) and strain at break, which decreased by 7.0-fold and 1.6-fold, respectively, consistent with the degradation-induced increase in porosity (fig. S5). Importantly, despite this gradual reduction in mechanical strength, the membrane retained sufficient structural integrity to maintain its architecture throughout the 7-day cell culture period without structural collapse.

### Formation of a lung fibroblast–derived ECM layer on the alveoli-mimetic PLGA membrane

To reconstruct the interstitial microenvironment of the alveolar niche, primary lung fibroblasts were cultured on the apical surface of the alveoli-mimetic PLGA membrane to generate a fibroblast-derived ECM layer. Using FITC-labeled PLGA membranes (fig. S6), ECM deposition and membrane integrity were monitored to determine whether ECM formation compensates for progressive membrane degradation during cell culture (fig. S7). During the 7-day culture period, the membrane gradually degraded, leaving a central pore approximately 200 μm in diameter. Notably, lung fibroblasts maintained a continuous cellular layer even across regions where the membrane had degraded (Fig. 3A). Collagen I, a major structural component of the ECM, was detected throughout the fibroblast layer, including regions where the membrane had degraded (Fig. 3B) (*37*). SEM imaging from the basolateral side further confirmed that the fibroblast layer bridged the ∼200 μm-diameter opening (Fig. 3C).

**Figure 3.**
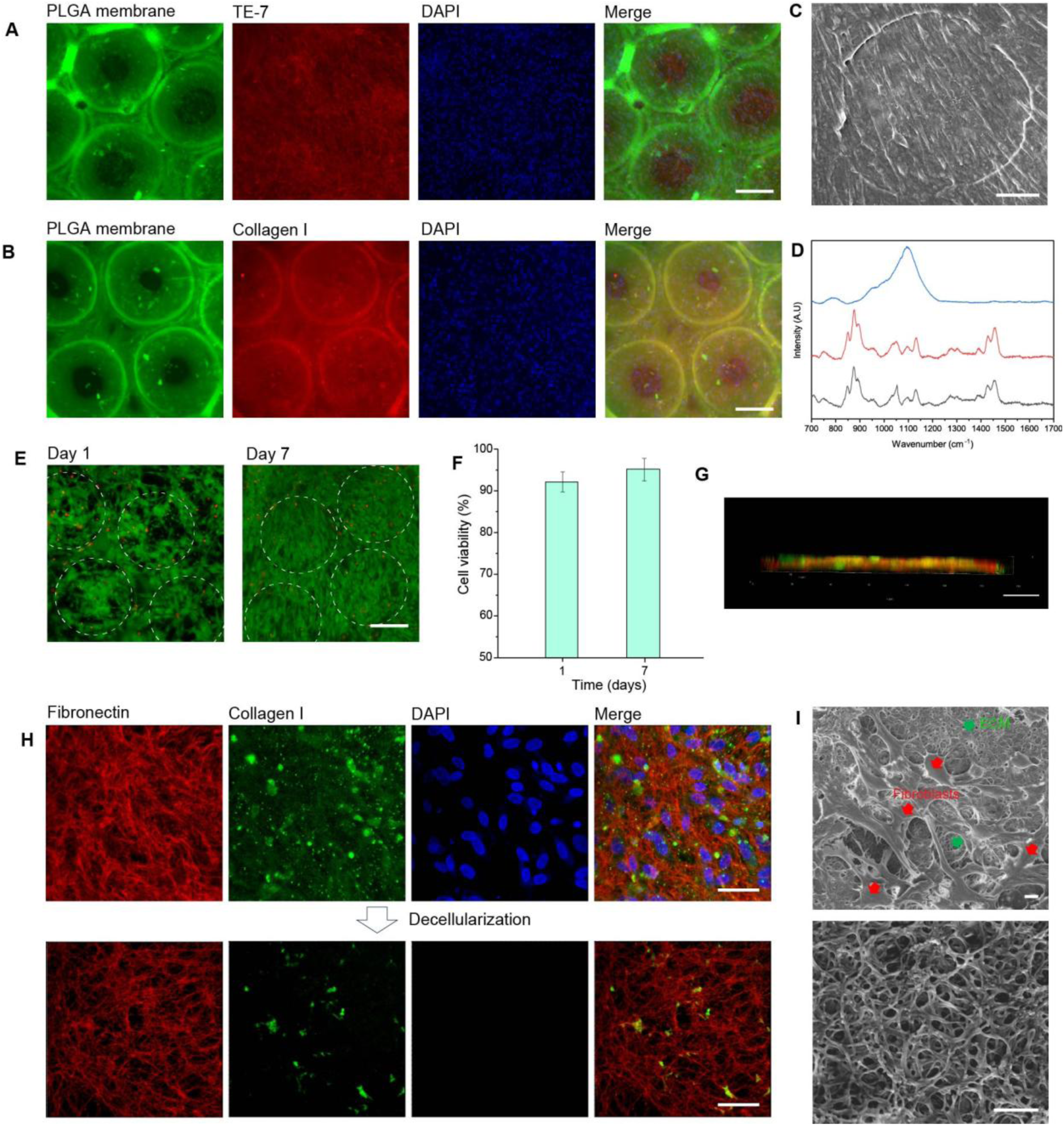
Formation of a stable lung fibroblast–derived ECM layer on alveoli-mimetic PLGA membranes. **(A)** Immunofluorescence image of lung fibroblasts cultured for 7 days on FITC-labeled alveoli-mimetic PLGA membranes, stained with TE-7 (red). Scale bar: 200 μm. (**B)** Immunofluorescence image of lung fibroblasts cultured for 7 days on FITC-labeled alveoli-mimetic PLGA membranes, stained with Collagen I (red). Scale bar: 200 μm. **(C)** SEM image of the lung fibroblast layer cultured for 7 days on the PLGA membrane, viewed from the basolateral side. Scale bar: 50 μm. **(D)** Raman spectra correspond to the cell-only region (blue), membrane-retained region (red), and pure PLGA (black). **(E)** Live/dead staining images of lung fibroblasts cultured on the chip at day 1 and day 7. Dashed circles indicate the alveolar structures. Scale bar: 200 μm. **(F)** Quantification of cell viability of lung fibroblasts cultured on the chip at day 1 and day 7 (n = 4). **(G)** Side-view immunofluorescence image of lung fibroblasts cultured for 7 days on the chip, stained for fibronectin (red) and collagen I (green). Scale bar: 30 μm. **(H)** Immunofluorescence images of the 7-day cultured lung fibroblast layer on the PLGA membrane, stained for fibronectin (red), collagen I (green), and DAPI (blue) (top). Following decellularization (bottom), cellular components were removed while the fibroblast-derived ECM network was preserved. Scale bars: 50 μm. **(I)** Low-magnification SEM image showing lung fibroblasts and the fibroblast-derived ECM cultured for 7 days on the chip (top), and a high-magnification SEM image highlighting the ultrastructure of the ECM network (bottom). Scale bars: 10 μm.

Chemical degradation of the PLGA membrane was further analyzed by Raman spectroscopy. After fibroblast culture, regions where the membrane had completely degraded and regions where residual membrane remained were selectively examined (Fig. S8). The regions with remaining membrane exhibited a characteristic PLGA peak at 1087 cm⁻¹, corresponding to ester bond stretching, consistent with the PLGA reference (Fig. 3D) (*38*). In contrast, this peak was absent in regions where only the cell-derived layer remained, confirming complete degradation of the PLGA membrane.

The viability of lung fibroblasts cultured on the PLGA membrane was assessed by live/dead staining (Fig. 3E). Throughout the 7-day culture period on the chip, fibroblasts maintained high viability, with more than 90% of cells remaining viable (Fig. 3F).

Fibronectin, another major ECM component that regulates cell adhesion and matrix organization (*37, 39*), was detected together with collagen I in fibroblasts cultured on the chip (Fig. 3, G and H). Fibronectin plays a critical role in ECM assembly by directing the deposition and organization of collagen fibrils within the matrix (*40*). Following decellularization, cellular components and collagen I were largely removed, whereas fibronectin remained on the membrane surface, indicating preservation of the fibroblast-derived ECM network (Fig. 3H). SEM imaging after decellularization further revealed that the remaining ECM formed a nanofibrous architecture on the membrane surface (Fig. 3I). These results demonstrate that the PLGA membrane was effectively replaced by a fibroblast-derived ECM layer.

To determine whether the fibroblast-derived ECM layer is required to support epithelial integrity after membrane degradation, epithelial cells were cultured on the chip in the absence of lung fibroblasts. Under these conditions, the epithelial layer failed to maintain structural continuity, and large openings formed at sites where the membrane had degraded (fig. S9). These results indicate that the fibroblast layer functions as a structural support that stabilizes the epithelial layer after membrane degradation.

### Formation of a functional epithelial barrier on the membrane-free alveolar chip

To recapitulate the alveolar microenvironment, human alveolar epithelial cells (A549 cells), which exhibit characteristics of type II alveolar epithelial cells (*41*), were used (fig. S10). Figure 4A illustrates the experimental workflow for establishing epithelial–fibroblast co-culture on the chip. Lung fibroblasts were first seeded on the membrane and cultured until a confluent monolayer formed. Subsequently, A549 epithelial cells were seeded on top of the fibroblast layer to form an epithelial monolayer. On day 5, the apical medium was removed to initiate ALI culture. During epithelial–fibroblast co-culture on the chip, the PLGA membrane progressively degraded over the 7-day culture period. As a control, fibroblasts were cultured on the basolateral side of the PCL membrane, which was considered non-degradable under the experimental conditions, whereas epithelial cells were seeded on the apical side (figs. S2, D to F). Despite membrane degradation, both epithelial and fibroblast cells formed stable and continuous cellular layers on the chip (Fig. 4, B and C, and fig. S11). Cross-sectional SEM images further confirmed that, in regions where the membrane had completely degraded, the fibroblast layer spanned the opening and acted as a structural support for the overlying epithelial layer (Fig. 4D). In regions where the membrane remained, the residual membrane exhibited a very thin, highly porous structure with an interconnected pore network. This architecture facilitates gas exchange and nutrient transport under ALI culture conditions. Side-view immunofluorescence imaging further confirmed the formation of an epithelial layer on top of the fibroblast layer on the chip (Fig. 4E).

**Figure 4.**
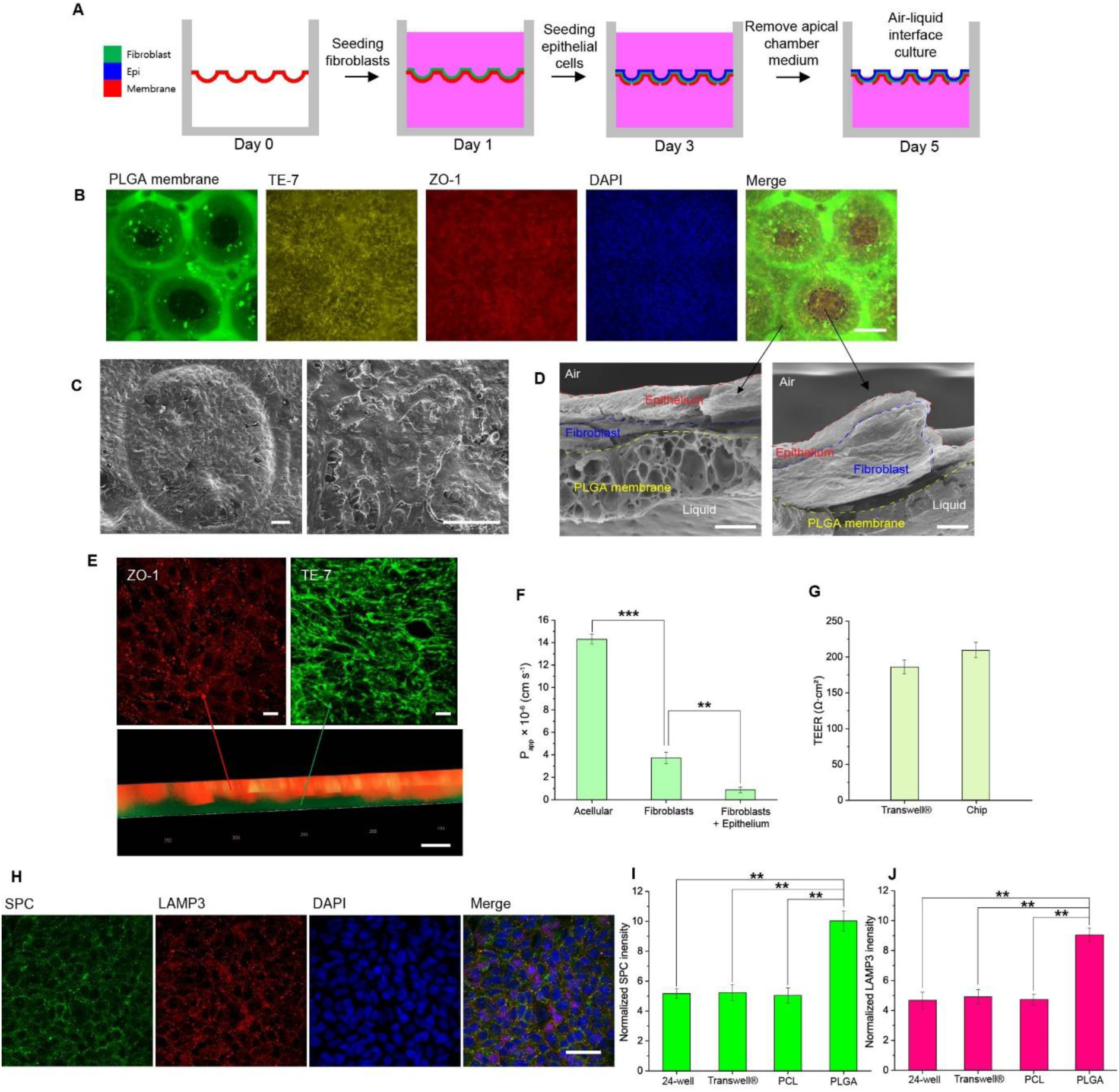
Epithelial–fibroblast co-culture under ALI conditions establishes a functional alveolar barrier. **(A)** Schematic illustration of the co-culture procedure for establishing the alveolar epithelial–fibroblast model on the alveoli-mimetic PLGA membrane. Lung fibroblasts were first seeded on the membrane (day 1), followed by seeding of A549 epithelial cells (day 3). On day 5, the apical medium was removed to initiate ALI culture. During the culture period, the PLGA membrane gradually degraded. **(B)** Immunofluorescence images of lung fibroblasts cultured on the PLGA membrane for 7 days and A549 cells cultured on top of the fibroblast layer for 5 days, including 2 days under ALI conditions. Lung fibroblasts were stained with TE-7, and A549 cells were stained with ZO-1. Scale bar: 200 μm. **(C)** SEM images of lung fibroblasts and A549 cells co-cultured on the chip for 7 days. The left panel shows a low-magnification top-view image of the co-culture, while the right panel shows a high-magnification image of A549 cells. Scale bars: 50 μm. **(D)** Cross-sectional SEM images of lung fibroblasts and A549 cells co-cultured on the chip for 7 days. The left panel shows a region where the membrane remains after partial degradation, while the right panel shows a region where the membrane has fully degraded, leaving only the cellular layer. Scale bars: 5 μm. **(E)** Immunofluorescence images of lung fibroblasts and A549 cells co-cultured on the chip, stained with TE-7 (green) and ZO-1 (red). The bottom panel shows a cross-sectional view highlighting the epithelial and fibroblast layers. Scale bars: 20 μm. **(F)** Apparent permeability (P_app_) of alveoli-mimetic PLGA membranes after 7 days of degradation, measured for acellular membranes, membranes cultured with lung fibroblasts, and membranes co-cultured with lung fibroblasts and A549 cells on the chip (n = 4) **(G)** Transepithelial electrical resistance (TEER) of lung fibroblasts and A549 cells co-cultured for 7 days on Transwell® membranes and on the chip. **(H)** Immunofluorescence images of A549 cells cultured on a fibroblast layer on the chip under ALI conditions for 2 days, stained for SPC, LAMP3, and DAPI. Scale bar: 50 μm. Quantification of SPC **(I)** and LAMP3 **(J)** fluorescence intensity in A549 cells cultured under four different conditions (24-well plate, Transwell®, PCL membrane, and PLGA membrane) (n = 4).

Barrier function was evaluated by measuring the apparent permeability across the cellular layers. The formation of a fibroblast layer reduced permeability by 1.3-fold compared with the acellular membrane. When epithelial cells were co-cultured with fibroblasts to form a double cellular layer, permeability decreased further by 2.1-fold (Fig. 4F). These results indicate that the epithelial layer functions as an effective protective barrier (*42*).

The TEER values of the epithelial–fibroblast double layer formed on the chip were comparable to those measured in Transwell® systems (Fig. 4G). Specifically, the epithelial–fibroblast co-culture exhibited TEER values of 204.4 Ω·cm² on the chip and 190.1 Ω·cm² in the Transwell® system. Notably, these values are substantially higher than previously reported TEER values for A549 epithelial monocultures (∼136 Ω·cm²) (*43*), indicating the formation of a robust epithelial barrier in the co-culture system. These results suggest that stable cellular barriers can be established on the chip even after the degradation and disappearance of the PLGA membrane. Together, these findings demonstrate that the fibroblast-derived ECM layer supports the formation of a functional epithelial barrier on the membrane-free chip.

To investigate the effect of direct fibroblast–epithelial cell contact on surfactant production, A549 epithelial cells were cultured under four different conditions: (i) on a fibroblast layer enabling direct contact (PLGA chip), (ii) on a 24-well plate (epithelial monoculture), (iii) on a Transwell® insert under ALI conditions (epithelial monoculture), and (iv) on a PCL membrane with fibroblasts cultured on the opposite side (fig. S2, D and E), preventing direct contact. Compared to conditions without fibroblasts or without direct cell–cell contact (fig. S12), direct contact between fibroblasts and epithelial cells significantly enhanced the expression of surfactant protein C (SPC), a canonical marker of alveolar type II epithelial cells responsible for pulmonary surfactant production (*44*), as well as lysosome-associated membrane protein 3 (LAMP3), which is associated with lamellar bodies involved in surfactant processing and secretion (Fig. 4, H to J) (*45, 46*). These results are consistent with previous findings that pulmonary fibroblasts promote epithelial surfactant production and suggest that the PLGA chip more closely recapitulates the physiological alveolar microenvironment (*47, 48*).

### PLGA mitigates stiffness-induced myofibroblast activation and epithelial injury

The apparent tensile modulus of the PLGA membrane was determined from its stress–strain curves and compared with that of Transwell® inserts (Fig. 5A). The PLGA membrane exhibited an apparent Young’s modulus of 8.55 MPa, which was approximately 47-fold lower than that of the Transwell® (*49*), indicating that the Transwell® substrate is substantially stiffer (Fig. 5B). Surface stiffness has been widely reported to regulate fibroblast differentiation, with stiffer substrates promoting the transition of fibroblasts into myofibroblasts (*50*), which play a key role in extracellular matrix remodeling and fibrotic responses in the alveolar microenvironment (*51*). To evaluate this effect, fibroblast differentiation on Transwell® inserts, and membrane-free regions was assessed by immunostaining for α-smooth muscle actin (α-SMA), a canonical marker of myofibroblast differentiation (Fig. 5C) (*52*). Fibroblasts cultured on the stiffer Transwell® substrate exhibited significantly elevated α-SMA expression, indicating enhanced myofibroblast differentiation (Fig. 5D). In contrast, fibroblasts in the membrane-free condition showed minimal α-SMA expression, indicating that the absence of a rigid substrate does not induce stiffness-driven myofibroblast differentiation.

**Figure 5.**
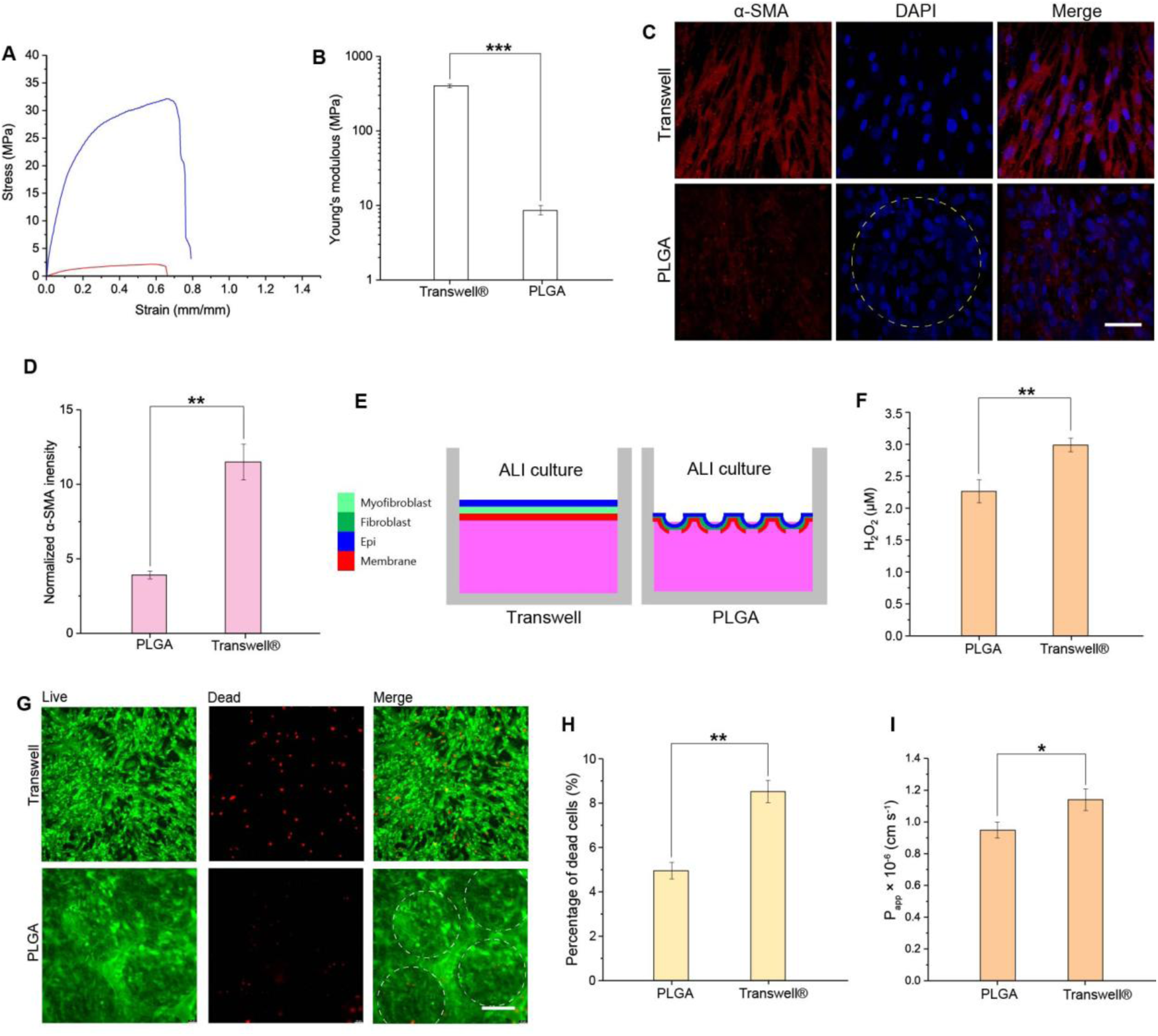
Rigid Transwell® substrates induce myofibroblast differentiation, oxidative stress, and cell death compared to biodegradable PLGA membranes. **(A)** Tensile stress–strain curves of the Transwell® membrane (blue) and the porous PLGA membrane prior to degradation (red). **(B)** Young’s modulus of the Transwell® membrane and the porous PLGA membrane prior to degradation (n = 4). **(C)** Immunofluorescence images of α-SMA expression in fibroblasts cultured on Transwell® and PLGA membranes after 7 days. The yellow dashed circles indicate regions where the PLGA membrane has degraded and is no longer present. Scale bar: 50 μm. **(D)** Quantification of α-SMA fluorescence intensity in fibroblasts cultured on Transwell® and PLGA membranes after 7 days (n = 4). **(E)** Schematic illustration of co-culture of fibroblasts and epithelial cells on Transwell® and PLGA membranes. In the Transwell® system, fibroblasts undergo differentiation into myofibroblasts. **(F)** Reactive oxygen species (ROS) levels in co-culture on Transwell® and PLGA chips. Hydrogen peroxide (H₂O₂) concentration was quantified 24 h after medium exchange (n = 4). **(G)** Live/dead assay images of co-culture on Transwell® and PLGA chips. Dashed circles indicate the alveolar structures. Scale bar: 200 μm. **(H)** Quantification of the percentage of dead cells in co-culture on Transwell® and PLGA chips (n = 4). **(I)** Apparent permeability (P_app_) of PLGA membranes after 7 days of degradation and Transwell® membranes in co-culture with lung fibroblasts and A549 cells on the chip (n = 4).

To investigate the effect of fibroblast-to-myofibroblast differentiation on epithelial cells, fibroblasts and epithelial cells were co-cultured on Transwell® inserts and PLGA membranes for comparative analysis (Fig. 5E and fig. S13). Consistent with previous reports that activated myofibroblasts produce elevated levels of reactive oxygen species (ROS), including hydrogen peroxide (H₂O₂), which can induce epithelial cell apoptosis (*53*), ROS levels were significantly higher in the Transwell® co-culture system compared to the PLGA system (Fig. 5F). Correspondingly, live/dead staining revealed increased cell death in the Transwell® condition, while the PLGA system maintained higher cell viability (Fig. 5G). Quantitative analysis further confirmed a significantly higher percentage of dead cells in the Transwell® group (Fig. 5H). Consistent with the increased myofibroblast activation observed in the Transwell® system, cell permeability was elevated compared to the PLGA condition (Fig. 5I). This increase in permeability is in line with previous reports showing that fibrotic remodeling is associated with compromised epithelial barrier function and enhanced permeability (*54*). These results suggest that substrate-induced myofibroblast differentiation is associated with increased ROS production, epithelial cell death, and compromised barrier integrity. In contrast, the PLGA-based system mitigates these effects, likely by suppressing stiffness-driven myofibroblast activation. Collectively, these findings indicate that the rigid Transwell® substrate promotes a pro-fibrotic microenvironment, whereas the PLGA-based system more effectively preserves epithelial viability and barrier function, thereby maintaining a more physiologically relevant cellular state.

### Chemokine-driven monocyte migration across the alveolar barrier

To investigate whether the membrane-free alveoli-on-a-chip enables immune cell recruitment similar to that observed *in vivo*, we examined the transmigration of monocytes across the epithelial–fibroblast barrier. In the lung, circulating monocytes are recruited to sites of inflammation and migrate across the alveolar barrier in response to chemokines such as monocyte chemoattractant protein-1 (MCP-1) (*55, 56*).

Fig. 6A schematically illustrates the experimental concept. In the conventional Transwell® (fig. S13) and PCL system (fig. S2), THP-1 monocytes seeded in the apical compartment were unable to migrate through the epithelial–fibroblast layer. In contrast, the biodegradable PLGA membrane used in the alveoli-on-a-chip enabled monocyte transmigration across the cellular barrier. Quantification of monocyte migration confirmed this observation (Fig. 6B). In the presence of MCP-1 (20 ng/mL), a significantly higher fraction of THP-1 cells migrated across the epithelial–fibroblast layer on the chip compared with conditions without chemokine stimulation. In contrast, virtually no monocyte migration was observed across the Transwell® and PCL membrane. These results indicate that the membrane-free architecture of the chip provides a permissive microenvironment for immune cell transmigration. Immunofluorescence analysis further confirmed the identity of transmigrated cells. THP-1 cells collected from the basolateral chamber stained positive for the monocyte marker CD14 (Fig. 6C and fig. S14) (*57*), confirming that the cells that traversed the epithelial–fibroblast barrier retained their monocyte phenotype. Together, these results demonstrate that the membrane-free alveoli-on-a-chip enables chemokine-driven monocyte recruitment across the alveolar barrier, recapitulating a key immune process that occurs during pulmonary inflammation *in vivo*.

**Figure 6.**
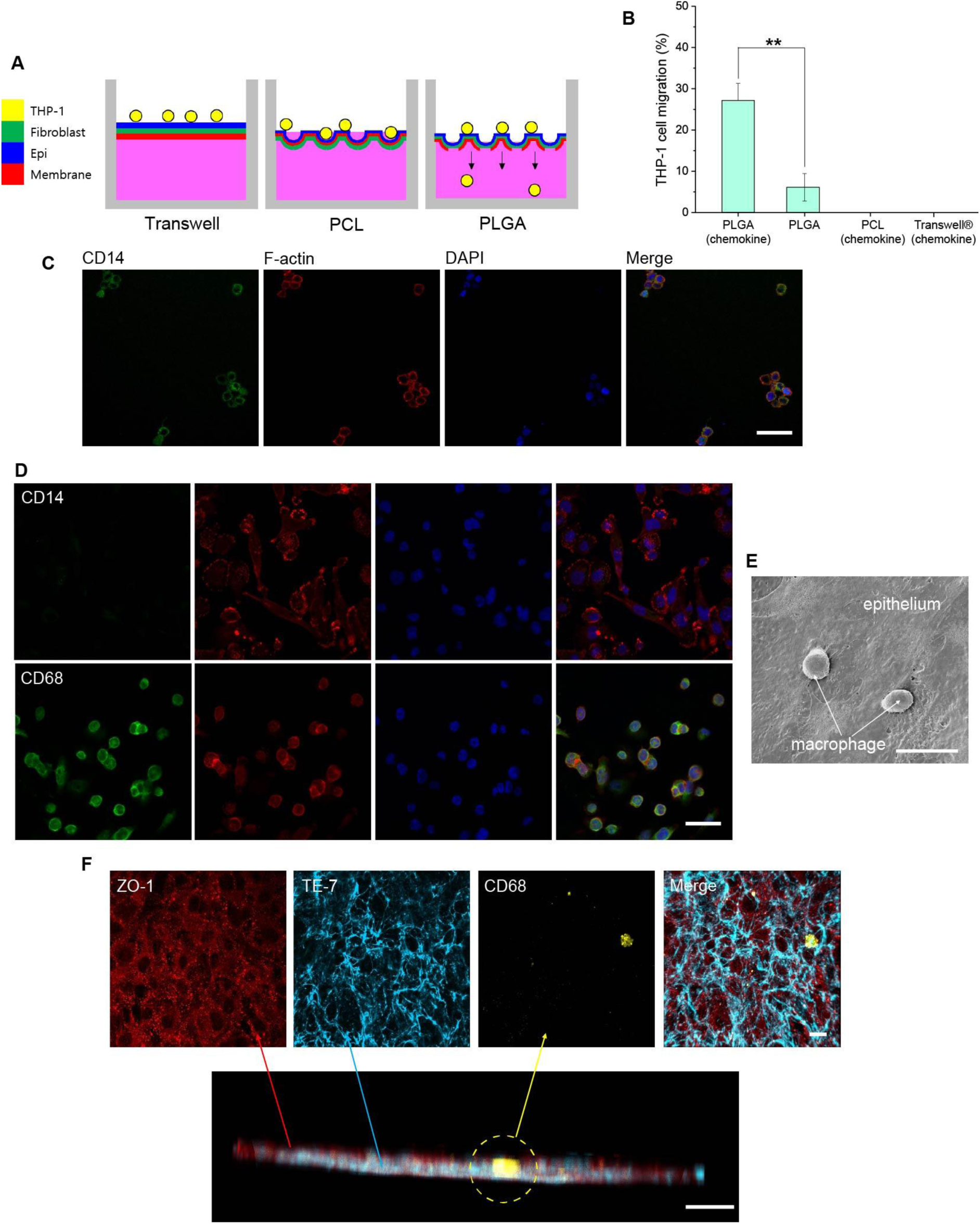
Monocyte migration and macrophage integration in the membrane-free alveoli-on-a-chip. **(A)** Schematic illustration of THP-1 monocyte migration across Transwell®, PCL, and PLGA membranes. Migration is restricted in Transwell® and PCL systems but enabled across the PLGA membrane. **(B)** Quantification of THP-1 cell migration after 24 h across epithelial–fibroblast co-cultures on the PLGA chip with or without MCP-1 (20 ng mL⁻¹), compared with Transwell® and PCL controls (n = 4). **(C)** Immunofluorescence images of THP-1 cells collected from the basolateral chamber after transmigrating through the fibroblast–epithelial layer. Cells were stained for CD14 (green) and F-actin (red), with nuclei counterstained with DAPI (blue). Scale bar: 50 μm. **(D)** Immunofluorescence images of THP-1 cells differentiated into macrophage-like cells following PMA treatment, stained for CD14 and F-actin (top) and CD68 and F-actin (bottom). Scale bar: 50 μm. **(E)** SEM image of macrophages on the chip under ALI conditions with fibroblast–epithelial co-culture. Scale bar: 30 μm. **(F)** Immunofluorescence images of lung fibroblasts, A549 cells, and THP-1–derived macrophages cultured on the alveoli-on-a-chip under ALI conditions. Fibroblasts were stained with TE-7, A549 cells were stained with ZO-1, macrophages were stained with CD68. The bottom panel shows a cross-sectional view highlighting the layered organization of the three cell types on the chip. Scale bars: 30 μm.

### Macrophage integration in the membrane-free alveoli-on-a-chip

To further recapitulate the immune component of the alveolar microenvironment, THP-1 monocytes were differentiated into macrophage-like cells using phorbol 12-myristate 13-acetate (PMA) (*58, 59*). Immunofluorescence staining confirmed successful differentiation of THP-1 monocytes into macrophage-like cells. While CD14 expression was minimal following differentiation, the cells exhibited strong expression of the macrophage marker CD68 along with pronounced F-actin cytoskeletal organization (Fig. 6D) (*60*).

The differentiated macrophage-like cells were subsequently introduced onto the epithelial surface of the alveoli-on-a-chip under ALI conditions. Macrophage-like cells adhered to the epithelial surface and remained viable within the chip microenvironment. Live–dead analysis demonstrated high overall cellular viability within the chip, with 93.9% of cells remaining viable under ALI culture conditions (Fig. 7, B and C), indicating that the chip environment supports immune cell integration and survival. SEM image further revealed macrophage morphology and interactions with the epithelial surface (Fig. 6E). The macrophages displayed characteristic rounded morphology and were positioned directly on the epithelial layer, resembling the localization of alveolar macrophages that reside on the epithelial surface of the alveolar lumen *in vivo*. Together, these results demonstrate that the membrane-free alveoli-on-a-chip supports macrophage integration under ALI conditions, enabling incorporation of a key immune component of the alveolar microenvironment.

**Figure 7.**
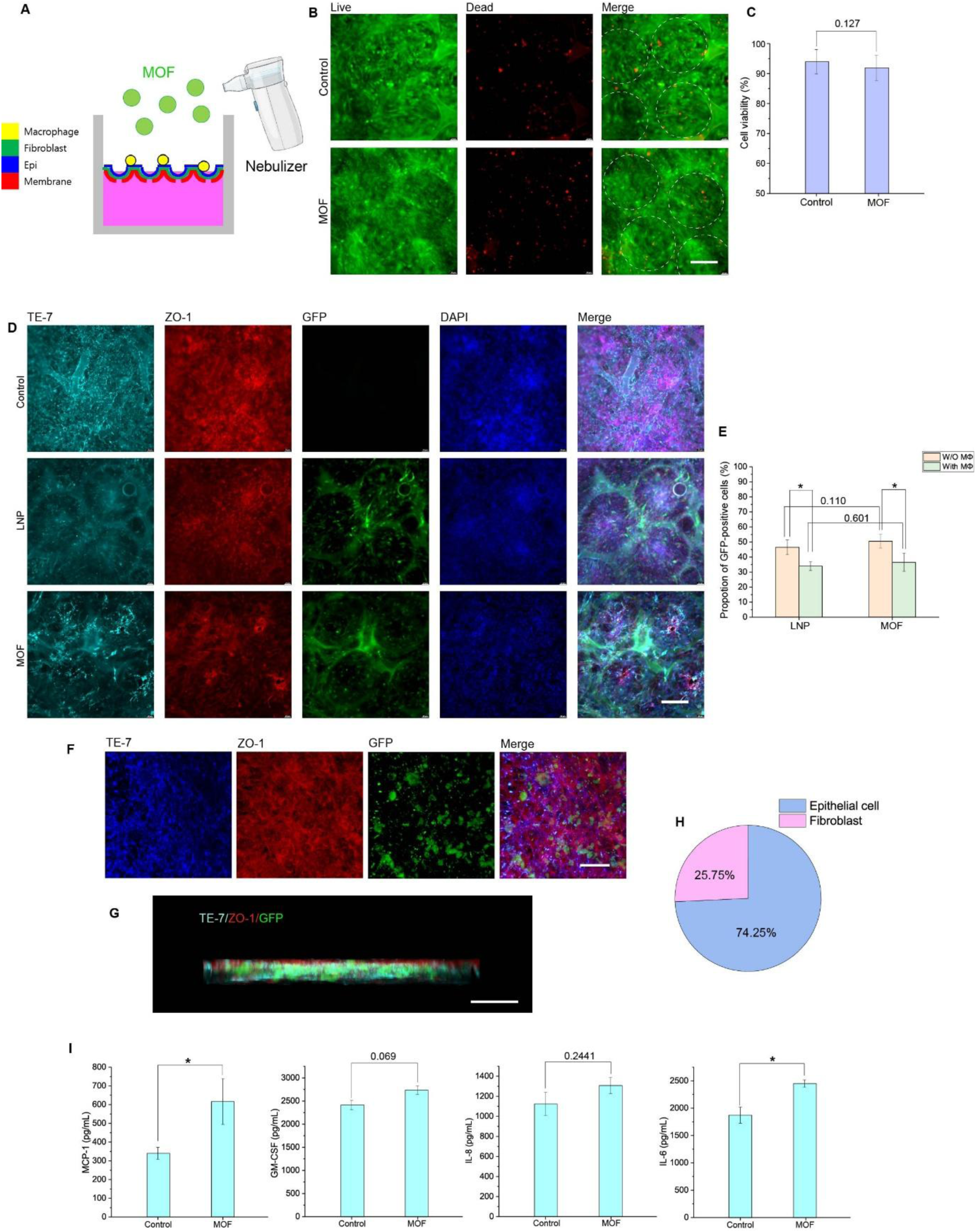
MOF nanoparticle aerosol delivery enables efficient gene transfection with minimal cytotoxicity in the alveoli-on-a-chip. **(A)** Schematic illustration of aerosol delivery of MOF nanoparticles to the alveoli-on-a-chip using a nebulizer. MOF nanoparticles (1 µg) were aerosolized and deposited onto the epithelial surface under air–liquid interface conditions and exposed to the chip for 48 h. **(B)** Live/dead fluorescence images of cells cultured on the chip after aerosol exposure to MOF nanoparticles (1 µg, 48 h) compared with untreated controls. Dashed circles indicate the alveolar structures. Scale bar: 200 μm. **(C)** Quantification of cell viability of cells cultured on the chip before and after MOF nanoparticle exposure (1 µg, 48 h) (n = 4). **(D)** Immunofluorescence images showing cellular responses in the alveoli-on-a-chip following nanoparticle delivery in the presence of macrophages. Fibroblasts were stained with TE-7, epithelial tight junctions of A549 cells were labeled with ZO-1, GFP indicates nanoparticle-mediated gene expression, and nuclei were counterstained with DAPI. Images correspond to control, LNP-treated (1 µg, 48 h), and MOF-treated (1 µg, 48 h) conditions. Scale bar: 200 μm. **(E)** Quantification of the proportion of GFP-positive cells following nanoparticle-mediated delivery in the presence or absence of macrophages (MΦ), comparing LNP and MOF nanoparticle formulations after exposure to nanoparticles (1 µg, 48 h) (n = 4). **(F)** Immunofluorescence images showing cellular responses in the macrophage-containing alveoli-on-a-chip following MOF treatment (1 µg, 48 h). Fibroblasts were stained with TE-7, epithelial tight junctions of A549 cells were labeled with ZO-1, GFP indicates nanoparticle-mediated gene expression, and nuclei were counterstained with DAPI. Scale bar: 50 μm. **(G)** Side-view immunofluorescence images showing cellular responses in the macrophage-containing alveoli-on-a-chip following MOF treatment (1 µg, 48 h). Scale bar: 30 μm. **(H)** Pie chart showing the proportion of MOF-transfected cells in the chip, indicating the relative distribution of fibroblasts and epithelial cells among GFP-positive cells. **(I)** Cytokine secretion profiles measured in the alveoli-on-a-chip 48 h after MOF nanoparticle exposure (1 µg), including MCP-1, GM-CSF, IL-8, and IL-6 (n = 3).

### Efficient aerosolized mRNA delivery on the membrane-free alveoli-on-a-chip

To evaluate the potential of the membrane-free alveoli-on-a-chip as a platform for pulmonary nanomedicine delivery, we examined aerosolized nanoparticle-mediated gene transfection on the chip. Metal–organic framework (MOF) nanoparticles, a class of porous crystalline materials composed of metal ions and organic linkers, have recently emerged as promising carriers for nucleic acid delivery due to their high cargo loading capacity, structural tunability, and physicochemical stability (*61*). These properties are particularly advantageous for pulmonary delivery, where nanoparticles must maintain structural integrity and functional activity during aerosolization (*62*). In this study, we employed a MOF nanoparticle formulation developed in our laboratory and compared its gene delivery performance with that of conventional lipid nanoparticles (LNPs), which represent the current standard platform for mRNA delivery (*30, 63, 64*). As illustrated in Fig. 7A, MOF nanoparticles carrying GFP-encoding mRNA were aerosolized using a nebulizer and deposited onto the epithelial surface of the chip under ALI conditions.

Cell viability following nanoparticle exposure was first evaluated to assess potential cytotoxicity. Live/dead staining revealed no detectable increase in cell death after exposure to aerosolized MOF nanoparticles compared with untreated controls (Fig. 7B). Quantitative analysis confirmed that overall cellular viability remained above 91.9% following 48 h of nanoparticle exposure (Fig. 7C), indicating that the nanoparticle delivery process did not compromise cellular viability within the alveoli-on-a-chip microenvironment.

We next examined the efficiency and cell-type specificity of nanoparticle-mediated gene delivery in the multicellular alveolar model. To first assess cell-type–dependent transfection efficiency, epithelial cells and lung fibroblasts cultured as monocellular layers were exposed to MOF and LNP formulations for 48 h (fig. S15). Under these conditions, LNP treatment resulted in detectable GFP expression primarily in fibroblasts, whereas little or no transfection was observed in epithelial cells. In contrast, MOF nanoparticles mediated gene delivery in both epithelial cells and fibroblasts.

Based on these results, nanoparticle-mediated gene delivery was further evaluated in the multicellular alveoli-on-a-chip model. Immunofluorescence imaging demonstrated GFP expression in both epithelial and fibroblast layers following aerosol delivery (Fig. 7D). In the case of LNP treatment, GFP signals were predominantly detected in fibroblasts beneath the epithelial layer, suggesting that LNP particles preferentially crossed the epithelial barrier and transfected the underlying fibroblast layer. In contrast, MOF nanoparticles mediated gene delivery in both epithelial cells and fibroblasts within the chip microenvironment. Quantification of GFP-positive cells indicated that MOF nanoparticles achieved a higher overall transfection rate compared with LNP formulations, although the difference was not statistically significant (Fig. 7E).

To further investigate the influence of immune cells on nanoparticle-mediated gene delivery, we compared transfection efficiencies in the absence (fig. S16) and presence of macrophages (Fig. 7D). When macrophages were introduced into the chip microenvironment, the overall transfection efficiency decreased for both nanoparticle formulations. Specifically, MOF-mediated transfection decreased from 50.5% to 36.5%, whereas LNP-mediated transfection decreased from 46.5% to 34.0%. This reduction is likely attributable to the phagocytic activity of macrophages, which can internalize and sequester nanoparticles before they reach target epithelial cells, consistent with the well-known role of alveolar macrophages as the first-line defense system that clears inhaled particulates from the alveolar space (*17, 19*).

Alveolar surfactant is a lipid-rich interfacial film composed primarily of phospholipids and surfactant proteins that coats the alveolar air–liquid interface (*65, 66*). Although surfactant deposition was confirmed in the chip under ALI culture conditions (Fig. 4H), both MOF and lipid nanoparticles were able to traverse this surfactant layer and successfully mediate gene delivery to the underlying epithelial and fibroblast compartments. Together, these findings demonstrate that MOF nanoparticles can effectively deliver functional mRNA to multiple cell types within the membrane-free alveoli-on-a-chip.

To further determine which cell populations were transfected within the chip microenvironment, immunofluorescence staining was performed for fibroblasts (TE-7), epithelial tight junctions (ZO-1), and GFP expression following MOF treatment (Fig. 7F). The results revealed that the majority of GFP-positive cells were localized within the epithelial layer, indicating that epithelial cells were the primary targets of MOF-mediated gene delivery. However, a subset of GFP-positive fibroblasts was also observed beneath the epithelial layer. Cross-sectional imaging further confirmed the spatial distribution of transfected cells across the alveolar tissue structure (Fig. 7G). Quantitative analysis demonstrated that 74.25% of the transfected cells were epithelial cells, whereas 25.75% were fibroblasts (Fig. 7H). These findings suggest that while MOF nanoparticles predominantly transfect alveolar epithelial cells, they are also capable of penetrating the epithelial barrier and delivering genetic material to fibroblasts within the alveolar interstitium, highlighting their potential to influence both epithelial and interstitial cellular compartments of the lung microenvironment.

To assess inflammatory responses induced by nanoparticle exposure, cytokine secretion profiles were measured in the chip microenvironment following aerosol delivery. After 48 h of exposure to MOF nanoparticles, a trend toward increased cytokine secretion was observed. Among the cytokines analyzed, MCP-1 and IL-6 levels were significantly elevated compared with untreated controls, whereas GM-CSF and IL-8 showed no statistically significant changes (Fig. 7I). These results suggest that MOF nanoparticle exposure induces a moderate inflammatory response in the multicellular alveolar model while maintaining overall cellular viability and barrier integrity (*67*).

Together, these findings demonstrate that aerosolized MOF nanoparticles enable efficient gene delivery in the membrane-free alveoli-on-a-chip while maintaining high cellular viability and only modest inflammatory responses. These results establish the membrane-free alveoli-on-a-chip as a physiologically relevant platform for evaluating inhaled nanomedicine delivery and gene therapy strategies.

## Discussion and Conclusion

In this study, we developed a membrane-free alveoli-on-a-chip enabled by a biodegradable PLGA scaffold that reconstructs key structural and functional features of the human alveolar microenvironment. By allowing the synthetic membrane to gradually degrade and be replaced by fibroblast-derived extracellular matrix (ECM), the system transitions from an artificial scaffold to a biologically generated interstitial layer, more closely resembling the native alveolar architecture.

A central feature of this platform is the dynamic reconstruction of the interstitial microenvironment. The porous PLGA membrane initially provides mechanical support and an alveolus-mimetic dome structure, while progressive degradation enables lung fibroblasts to deposit ECM components such as collagen and fibronectin. This process results in a continuous, cell-derived matrix that maintains structural integrity and supports epithelial organization even after the disappearance of the synthetic membrane. The fibroblast-derived ECM was essential for maintaining epithelial continuity, as epithelial cells failed to sustain a stable layer in the absence of fibroblasts during membrane degradation.

Beyond structural support, the reconstructed interstitial environment plays a critical role in regulating epithelial phenotype. Direct contact between fibroblasts and epithelial cells significantly enhanced the expression of surfactant-related markers, including surfactant protein C (SPC) and LAMP3, compared to monoculture or non-contact conditions. These findings indicate that both ECM-mediated support and direct epithelial–fibroblast interactions contribute to epithelial maturation and surfactant-associated functions, highlighting the importance of recreating the interstitial niche for achieving physiologically relevant epithelial behavior.

Importantly, our results further demonstrate that interstitial mechanics critically regulate cellular responses within the alveolar microenvironment. We found that conventional rigid Transwell® substrates promote fibroblast-to-myofibroblast differentiation, as evidenced by elevated α-SMA expression. This phenotypic transition was associated with increased reactive oxygen species (ROS) production, enhanced epithelial cell death, and compromised barrier function. In contrast, the membrane-free PLGA system, particularly in regions where the rigid substrate is absent, suppressed stiffness-driven myofibroblast activation and preserved epithelial viability and barrier integrity. These findings reveal that widely used membrane-based systems can introduce non-physiological mechanical cues that drive pathological responses, and underscore the importance of recapitulating a mechanically relevant interstitial microenvironment.

The membrane-free architecture also enabled physiological cellular processes that are difficult to reproduce in conventional models. Chemokine-driven monocyte migration across the epithelial–fibroblast barrier was observed in response to MCP-1 stimulation, whereas migration was restricted in Transwell® and non-degradable PCL systems. This suggests that the absence of a permanent synthetic barrier provides a more permissive and physiologically relevant microenvironment for immune cell trafficking. In addition, the platform supported the integration and survival of macrophages on the epithelial surface under air–liquid interface conditions, further enhancing the physiological relevance of the model.

Beyond modeling alveolar biology, the membrane-free alveoli-on-a-chip provides a robust platform for evaluating inhaled nanomedicine delivery. Aerosolized MOF nanoparticles successfully mediated mRNA delivery to both epithelial and fibroblast layers while maintaining high cellular viability and inducing only moderate inflammatory responses. Compared with lipid nanoparticles, MOF nanoparticles demonstrated broader cellular transfection within the multicellular alveolar model, highlighting their potential as effective carriers for pulmonary gene delivery.

Overall, this membrane-free alveoli-on-a-chip bridges a critical gap between conventional membrane-based *in vitro* systems and the native alveolar microenvironment. By integrating dynamic ECM remodeling, direct epithelial–fibroblast interaction, physiologically relevant mechanical cues, immune cell trafficking, and nanoparticle-mediated gene delivery, this platform provides a versatile tool for studying lung biology and disease mechanisms. In particular, the ability to simultaneously promote epithelial maturation while suppressing stiffness-driven pathological signaling underscores the potential of this system for investigating fibrosis-related processes and for developing more effective inhaled therapeutic strategies.

## Materials and methods

### Fabrication process of curved microstructure arrays

The fabrication process, as shown in fig. S1, begins with a polystyrene (PS) microsphere suspension (10 wt%, Sigma-Aldrich), which is redispersed in a 1:1 mixture of deionized water and ethanol. The suspension is deposited at the water–air interface in an oxygen-plasma-treated PDMS reservoir using a tilted blade to enhance wettability and ensure uniform spreading. Driven by the Marangoni effect caused by surface tension gradients between ethanol and water, the PS microspheres disperse across the interface and self-assemble into a hexagonally close-packed monolayer through capillary interactions during solvent evaporation (A and B). After monolayer formation, the water is gradually replaced with a pre-polymer solution to form a hydrogel. The hydrogel is synthesized via free-radical polymerization of N-isopropylacrylamide (NIPAM) (Sigma-Aldrich, USA) with N,N′-methylenebis(acrylamide) (BIS) (Sigma-Aldrich, USA) as the crosslinker, followed by the addition of potassium persulfate (KPS) (Sigma-Aldrich, USA) and tetramethylethylenediamine (TEMED) (BioRad, USA) to initiate and accelerate the polymerization. The solution is introduced into the chamber containing the PS monolayer and allowed to polymerize under nitrogen for at least 1 hour (C and D). Subsequent hydration causes hydrogel swelling, which increases the spacing between embedded PS microspheres (E and F). For mold fabrication, a 10:1 PDMS elastomer-to-curing-agent mixture (Dow SYLGAR 184 Silicone Elastomer kit, Ellsworth Adhesives, USA) is poured onto the hydrogel assembly and cured at room temperature for 24 hours. After curing, the PDMS layer is peeled off, transferring the PS microspheres onto the PDMS surface (G and H). The microspheres are then removed using acetone or toluene, producing a concave PDMS mold. This mold is used to fabricate convex microstructures by casting blue silicone rubber (LET’S RESIN, Hong Kong), which is cured overnight (I and J). The cured layer was subsequently demolded to obtain the final convex microstructure array. Finally, a thin layer of pre-polymer solution was spin-coated onto the mold to fabricate a thin polymeric membrane (K and L).

### PLGA membrane fabrication

PLGA (poly(D,L-lactide-co-glycolide), lactide:glycolide = 50:50; Sigma-Aldrich, P2191) pellets were used to fabricate porous membranes. A 10 wt% PLGA solution was prepared by dissolving PLGA in acetone at 45 °C under magnetic stirring for 3 h. Camphene (100 wt% relative to PLGA; Sigma-Aldrich, 456055) was then added to the PLGA solution and further stirred at 45 °C for 1 h to obtain a homogeneous PLGA–camphene mixture. The solution was subsequently cooled to room temperature prior to membrane fabrication. Porous PLGA membranes were fabricated by spin-coating the PLGA–camphene solution onto an alveoli-shaped mold at 400 rpm for 3 s. The coated substrate was immediately immersed in distilled water at room temperature for 10 min to induce solidification of the PLGA and camphene phases. To generate a porous membrane structure, camphene was removed by freeze-drying for 3 h.

### Membrane characterization

The surface morphology and porous structure of the PLGA membranes were examined using scanning electron microscopy (SEM; Quanta 600 FEG, FEI Company, USA). For samples containing cultured cells, membranes were fixed in 2.5% glutaraldehyde for 1 h at room temperature. After three washes with phosphate-buffered saline (PBS), samples were post-fixed with 1% osmium tetroxide (OsO₄) for 1 h at 4 °C and subsequently dehydrated through a graded ethanol series (50–100%). Final dehydration was performed using hexamethyldisilazane (HMDS, Sigma Aldrich) for 15 min, followed by air drying prior to imaging.

The structural characteristics of the PLGA membranes were quantified from SEM images using ImageJ software. For each sample, five central pores were measured to determine the diameter of the alveolar openings, while fifty micropores were measured to quantify micropore size. Surface porosity was calculated by analyzing four SEM images per sample.

### PLGA membrane degradation assessment

PLGA membranes were incubated in phosphate-buffered saline (PBS) at 37 °C for predetermined time intervals to evaluate degradation behavior. After incubation, the membranes were gently rinsed with distilled water and subsequently freeze-dried for 3 h to remove residual moisture from the pores. The dry weight of the degraded membranes (Wₜ) was then measured. The degradation of the membranes was quantified by calculating the mass loss according to the following equation: Mass loss (%) = (W_0_−W_t_) / W_0_ ×100, where W_0_ represents the initial dry weight of the membrane and W_t_ represents the dry weight after degradation.

### Permeability measurement

Membrane permeability was evaluated using a FITC–dextran transport assay. Prior to the experiment, the culture medium was removed and the device was gently rinsed with PBS. Subsequently, 1 mL of phenol red–free culture medium was added to the basolateral chamber, while 0.5 mL of FITC–dextran solution (4 kDa; Sigma-Aldrich) was introduced into the apical chamber. The device was then incubated at 37 °C. Samples were collected from the basolateral compartment at 0, 10, 20, 30, and 40 min and transferred to a black 96-well plate. Fluorescence intensity was measured immediately after collection using a microplate reader (SpectraMax i3x) with excitation and emission wavelengths of 490 nm and 525 nm, respectively. The apparent permeability coefficient (P_app_, cm s⁻¹) was calculated using the following equation:

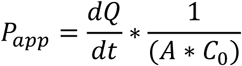

where dQ/dt is the transport rate of FITC–dextran across the membrane, A represents the effective membrane area (cm²), and C_0_ is the initial concentration of the fluorescent tracer in the apical chamber (mg mL⁻¹). The apparent permeability of the cell layer was calculated using the following equation (*23*).

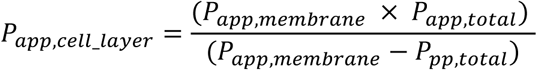

### Fourier transform infrared spectroscopy

The chemical composition of the PLGA membranes was characterized using FT-IR spectroscopy (Perkin Elmer Inc., USA). Spectra of camphene, PLGA membranes, and pure PLGA were acquired over a wavenumber range of 600–2000 cm⁻¹, with 32 scans collected for each sample.

### Mechanical characterization of PLGA membranes

The mechanical properties of PLGA membranes were evaluated by uniaxial tensile testing using a mechanical testing system (MTS Criterion® 42). Samples were stretched at a crosshead speed of 10 mm/min with a load capacity of 50 N while recording stress–strain responses. The Young’s modulus was determined from the linear region of the resulting stress–strain curves. Membrane thickness was measured from cross-sectional SEM images.

### Device assembly

To assemble the lung alveoli-on-a-chip device, the PLGA membrane was initially bonded to the apical PDMS chamber (surface area: 1.13 cm²) using uncured PDMS as an adhesive and cured overnight at 40 °C to achieve robust attachment. The apical chamber with the integrated membrane was subsequently aligned and bonded to the basolateral PDMS chamber using a liquid PDMS, followed by curing at 40 °C overnight to ensure complete and stable sealing.

### Cell culture

For cell culture, the devices were sterilized by UV irradiation for 3 h prior to use. Primary human lung fibroblasts (ATCC, PCS-201-013) were seeded into the apical chamber at a density of 4 × 10⁵ cells cm⁻². The fibroblasts were cultured in Fibroblast Basal Medium (ATCC, PCS-201-030) supplemented with the Fibroblast Growth Kit (ATCC, PCS-201-041) under liquid–liquid culture (LLC) conditions at 37 °C in a humidified atmosphere containing 5% CO₂ for up to 3 days until a confluent monolayer was formed. On day 3, A549 epithelial cells (ATCC, CCL-185) were seeded onto the fibroblast layer at a density of 3 × 10⁵ cells cm⁻² and maintained in a 1:1 mixture of Fibroblast Basal Medium and RPMI 1640. On day 5, the apical medium was removed to establish ALI culture conditions. Under ALI conditions, THP-1 monocytes or PMA-differentiated macrophages were seeded onto the epithelial layer at a density of 1 × 10⁵ cells cm⁻². The chip system was subsequently maintained under ALI conditions, and the culture medium was replaced every two days through the basolateral chamber.

### Fabrication of fluorescent PLGA membranes

Fluorescent PLGA membranes were fabricated by incorporating fluorescein isothiocyanate (c)-labeled PLGA particles (Mw 5 kDa, Nanosoft Polymers) into the PLGA solution described above at a mass ratio of 1:20 (fluorescent PLGA: non-fluorescent PLGA).

### Raman spectroscopy

Raman spectra were acquired using a confocal Raman microscope (LabRAM Soleil, Horiba). A 532 nm laser was employed as the excitation source with a maximum power of 29 mW. The laser beam was focused onto the sample using a 50× objective lens (NA = 0.60), resulting in an approximate spot size of ∼1.4 µm in diameter. Scattered light was collected in a backscattering configuration through the same objective and dispersed using a grating of 1800 grooves mm⁻¹. Spectra were acquired with an integration time of 1 s and 60 accumulations.

### Decellularization

To remove cellular components while preserving the extracellular matrix (ECM), samples were gently washed twice with PBS. A decellularization solution consisting of 20 mM ammonium hydroxide and 0.5% Triton X-100 in PBS was then added to fully cover the samples and incubated for 3 minutes at room temperature. Following decellularization, the samples were washed 3–5 times with PBS to remove residual cellular debris. For additional removal of nucleic acids, samples were optionally treated with DNase I (100 U mL⁻¹) for 10 minutes at room temperature, followed by three washes with PBS. For immunofluorescence analysis, the samples were subsequently fixed with 4% PFA for 10 minutes at room temperature prior to staining.

### Immunofluorescence microscopy

Cells cultured on the membranes were rinsed three times with phosphate-buffered saline (PBS) and fixed with 4% paraformaldehyde (Sigma-Aldrich) for 15 min at room temperature. After fixation, the membranes were carefully cut, and the cells were permeabilized with 0.1% Triton X-100 in PBS for 10 min at room temperature. The samples were then blocked with PBS containing 5% donkey serum for 45 min at room temperature. Primary antibodies (listed in Table S1) were diluted in blocking buffer and incubated with the samples overnight at 4 °C. After washing with PBS, the samples were incubated with the corresponding secondary antibodies for 1 h at room temperature. Nuclei were counterstained with DAPI following secondary antibody incubation. Fluorescence images were obtained using a Leica DMi8 fluorescence microscope or a Zeiss LSM 880 confocal microscope.

### Cell viability

Cell viability was evaluated using a Viability/Cytotoxicity Assay Kit (Biotium, USA). Samples were first gently rinsed with phosphate-buffered saline (PBS) and subsequently incubated in a staining solution containing 2 µM calcein AM and 4 µM ethidium homodimer III (EthD-III) diluted in culture medium. Staining was carried out at 37 °C for 20 min. After incubation, the samples were washed twice with PBS and transferred to fresh culture medium prior to imaging. Fluorescence images were obtained using a Leica DMi8 microscope with a 20× objective, and cell viability was quantified by analyzing four randomly selected fields per sample.

### Transepithelial electrical resistance

The barrier integrity of cell layers cultured on PLGA membranes was assessed by measuring transepithelial electrical resistance (TEER). Measurements were conducted using a chopstick electrode (MERSSTX01, Millipore, USA). The resistance of the cellular layer (R_cell_) was determined by subtracting the resistance of a blank device without cells from the total recorded resistance. TEER values (Ω·cm²) were calculated by multiplying R_cell_ (Ω) by the effective membrane area (cm²). For measurements under ALI conditions, 200 µL of culture medium was temporarily added to the apical chamber immediately before measurement to establish stable electrical contact.

### THP-1 cell migration assay

To induce chemotactic migration, the chemokine MCP-1 (PeproTech) was added to the basolateral chamber at a concentration of 20 ng mL⁻¹. THP-1 cells (2 × 10⁵ cells) were introduced into the apical chamber and allowed to migrate for 24 h under ALI conditions. After 24 h of incubation, cells that migrated to the basolateral chamber were collected and counted using a hemocytometer.

### THP-1 monocyte-to-macrophage differentiation

THP-1 monocytes (ATCC, TIB-202) were differentiated into macrophage-like cells by treatment with phorbol 12-myristate 13-acetate (PMA, 50 ng mL⁻¹) in RPMI 1640 medium for 24 h. After PMA stimulation, the medium was replaced with fresh PMA-free medium, and the cells were incubated for an additional 24 h to allow stabilization of the macrophage-like phenotype. After stabilization, the cells were detached using Accutase Cell Dissociation Solution (Thermo Fisher Scientific, A1110501).

### Nanoparticle formulation for mRNA delivery

DLin-MC3-DMA (MC3; MedKoo Biosciences), 1,2-dimyristoyl-rac-glycero-3-methoxy(poly(ethylene glycol) (DMG-PEG, Avanti Polar Lipids), and 1,2-distearoyl-sn-glycero-3-phosphocholine (DSPC, Avanti Polar Lipids) were used as received. Cholesterol, zinc nitrate hexahydrate, 2-methylimidazole, and linear polyethyleneimine (PEI, 20 kDa) were purchased from Sigma Aldrich. Ethanol (EtOH, 200 Proof) and citrate buffer (0.5 M, pH 4) were purchased from Fisher Scientifsic. EGFP mRNA (5moU) was purchased from Tri-Link BioTechnologies. Metal-organic framework (MOF) nanoparticles, ZIF-8 formulation, were synthesized following our previous work (*30*). Zinc nitrate hexahydrate (40 mM) and 2-methylimidazole (160 mM) precursor solutions were prepared in ethanol. EGFP mRNA was mixed with 2-methylimidazole solution, followed by the addition of linear PEI 20 kDa to a 9.3 N/P ratio. The mixture was incubated for 15 min at room temperature. Zinc nitrate solution was then added to initiate MOF formation, followed by incubation at room temperature for 2 h. The final MOF nanoparticles were collected by centrifugation, washed with ethanol, and stored at −80 °C until use. Benchmark lipid nanoparticles (LNPs) were prepared using a pipette-mixing method adapted from previously reported protocols (*68*). For the ethanol phase, lipids were dissolved in ethanol at a molar ratio of 50:10:38.5:1.5, composed of DLin-MC3-DMA, DSPC, cholesterol, and DMG-PEG 2000, respectively. The aqueous phase consisted of EGFP mRNA dissolved in 10 mM citrate buffer. Upon rapid mixing of the ethanol and aqueous phases by pipetting, LNPs self-assembled spontaneously via nanoprecipitation. The resulting LNP suspension was dialyzed against 1× phosphate-buffered saline (PBS) using a dialysis cassette for 2 h to remove residual ethanol. Purified LNPs were stored at 4 °C until further use.

### Hydrogen peroxide and multiplex cytokine analysis

Hydrogen peroxide (H₂O₂) levels were quantified using a Hydrogen Peroxide Assay Kit (Sigma-Aldrich, USA), and the levels of multiple cytokines were measured using a Multiplex Human Cytokine ELISA Kit (Anogen), both according to the manufacturers’ instructions.

### Quantification of Cell-Type–Specific Transfection

Cell-type–specific transfection efficiency was quantified using ImageJ software. GFP-positive cells were identified based on fluorescence signals. GFP signals colocalizing with the ZO-1–defined epithelial boundaries were classified as epithelial cell transfection, whereas GFP signals that did not overlap with ZO-1 staining were considered fibroblast transfection. For each sample, three representative images were analyzed, and a total of four independent samples were included in the analysis.

### Statistics

Statistical analyses were performed using Student’s *t*-test for comparisons between two groups. For experiments involving three or more groups, one-way or two-way analysis of variance (ANOVA) followed by Tukey’s post hoc test was applied. Differences were considered statistically significant when *p* < 0.05.

