## Supplemental File for "Membrane-Free Alveolus-on-a-Chip via Biodegradable Scaffold Recapitulates Interstitial Mechanics, Immune Trafficking, and Aerosolized mRNA Delivery"

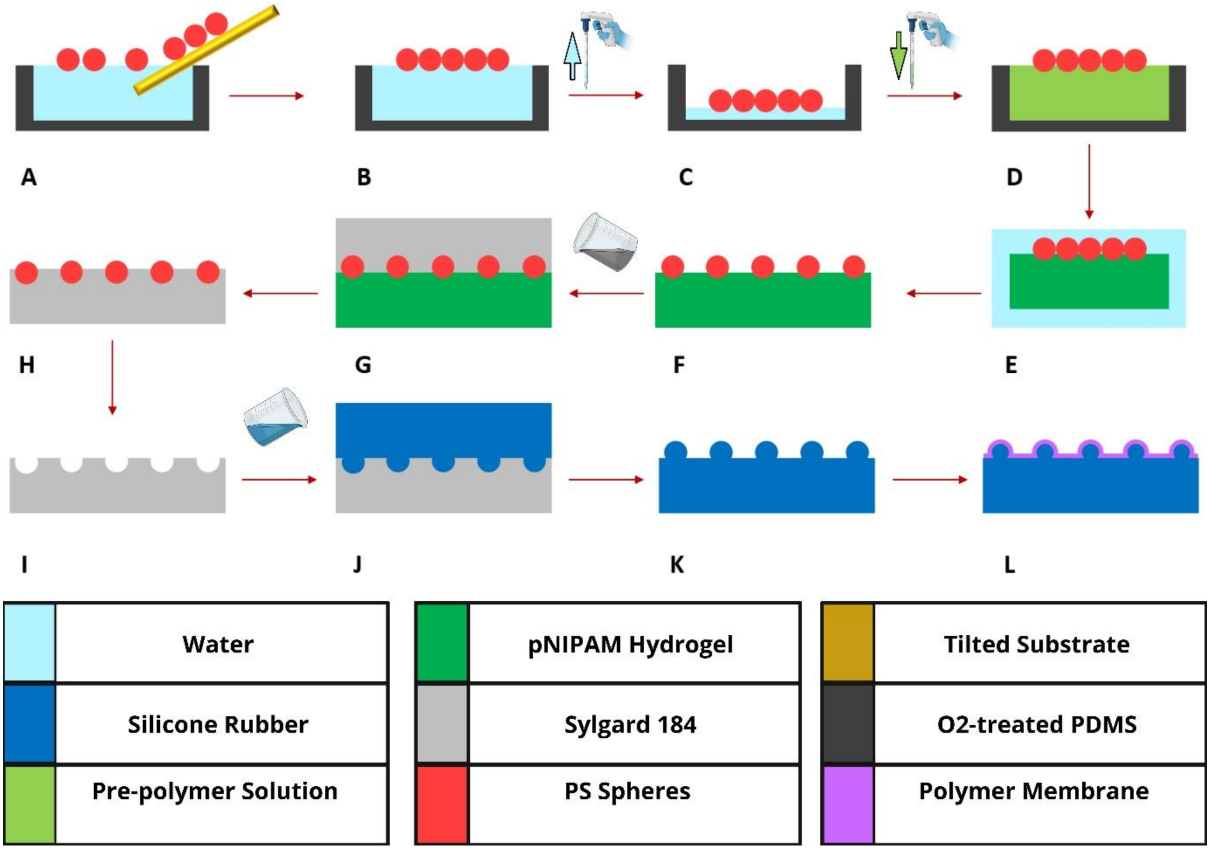


**Supplementary Figure 1.** Stepwise fabrication of alveoli-mimetic PLGA membranes using a PDMS mold.

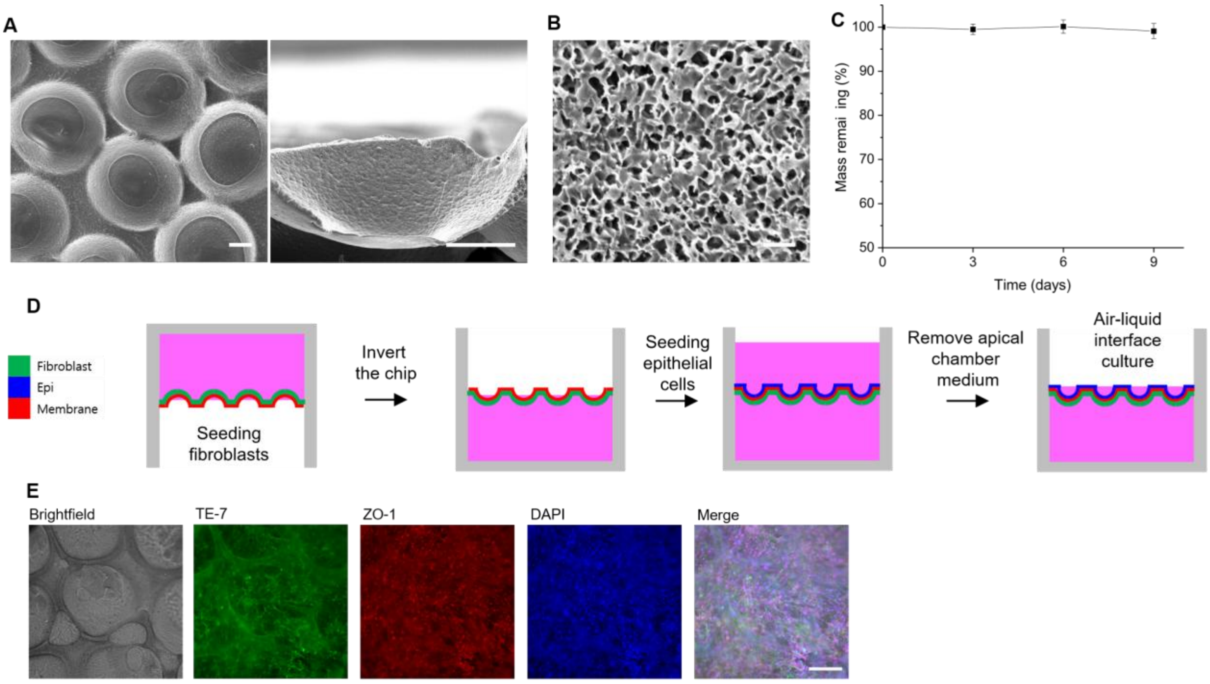


**Supplementary Figure 2. Structural characterization and cell culture of alveoli-mimetic PCL membrane-based chip (A)** Representative SEM images of alveoli-mimetic PCL membranes, showing the top view (left) and cross-sectional view (right). Scale bars: 100 μm. **(B)** Representative SEM image showing the microporous structure of the alveoli-mimetic PCL membrane. Scale bar: 10 μm. **(C)** Time-dependent mass loss of alveoli-mimetic PCL membranes during incubation in PBS at 37 °C (n = 4). **(D)** Schematic illustration of cell culture on the PCL membrane-based chip. Fibroblasts were first seeded on the basolateral side. After 3 h, once the fibroblasts adhered to the surface, the chip was inverted and epithelial cells were seeded on the apical side. From day 5, the apical medium was removed to initiate ALI culture. **(E)** Representative bright-field and fluorescence images of fibroblasts (basolateral) and A549 epithelial cells (apical) cultured on a PCL membrane under ALI conditions, stained for TE-7 and ZO-1, respectively. Scale bar: 200 μm.

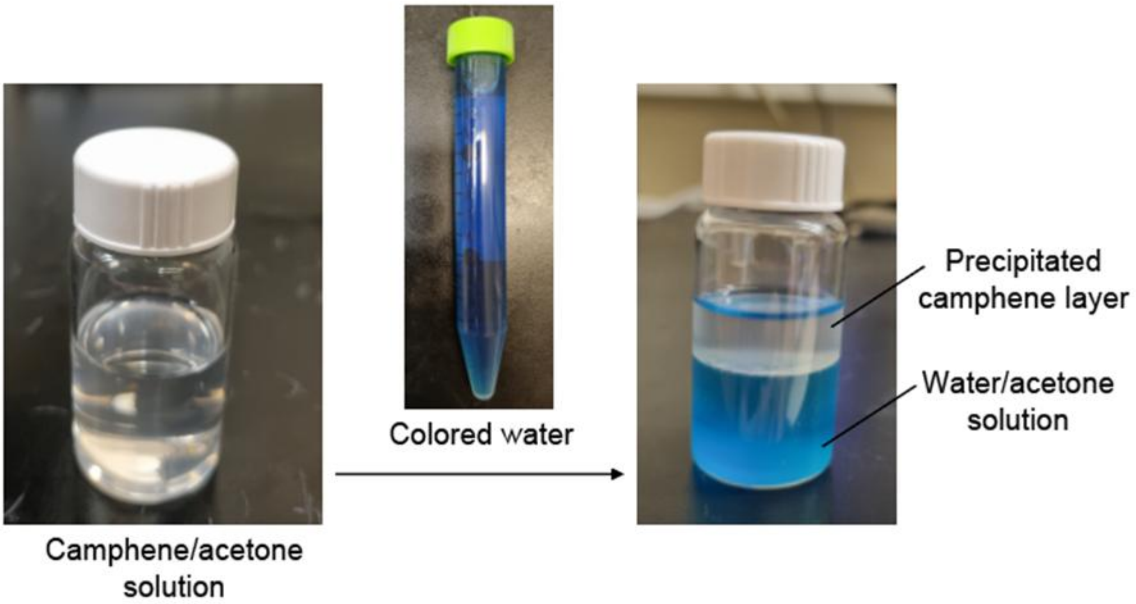


**Supplementary Figure 3.** Optical images showing the camphene/acetone solution (left) and the formation of a precipitated camphene layer on top of a water/acetone mixture after colored water was added to the camphene/acetone solution (right).

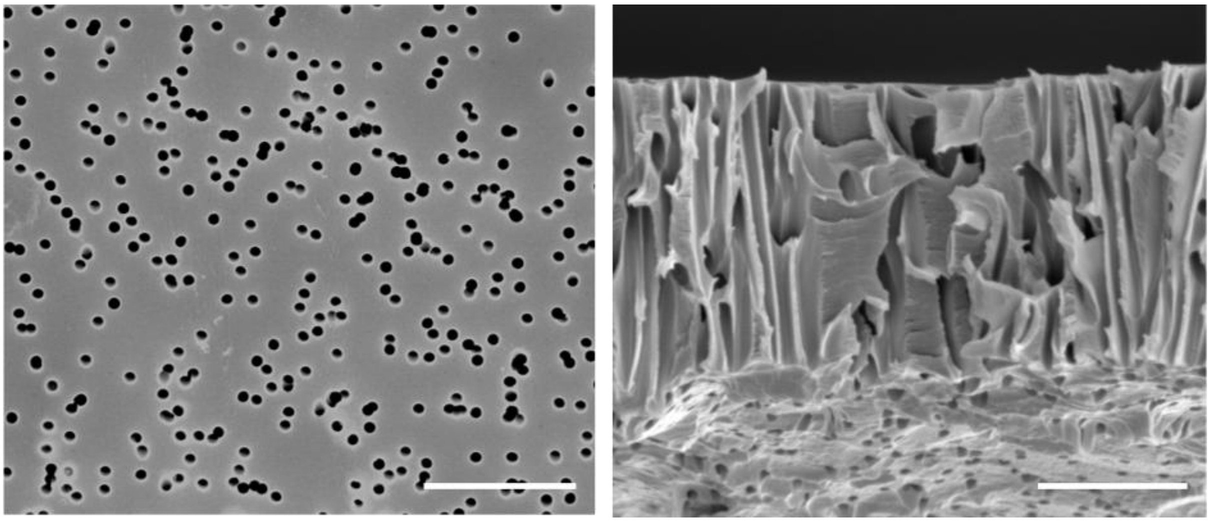

**Supplementary Figure 4.** SEM images of the 0.4 µm pore-sized Transwell® membrane showing the top view (left) and cross-sectional view (right). Scale bars: 5 µm.

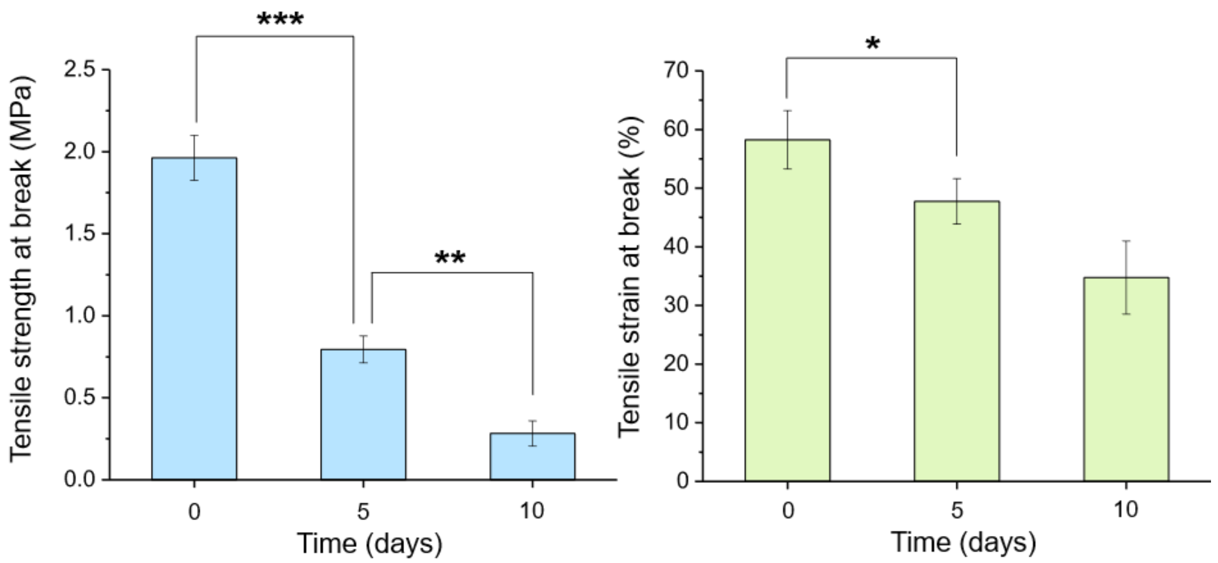


**Supplementary Figure 5** Tensile strength at break (left) and tensile strain at break (right) of alveoli-mimetic PLGA membranes after degradation in PBS at 37 °C for 0, 5, and 10 days (n = 4).

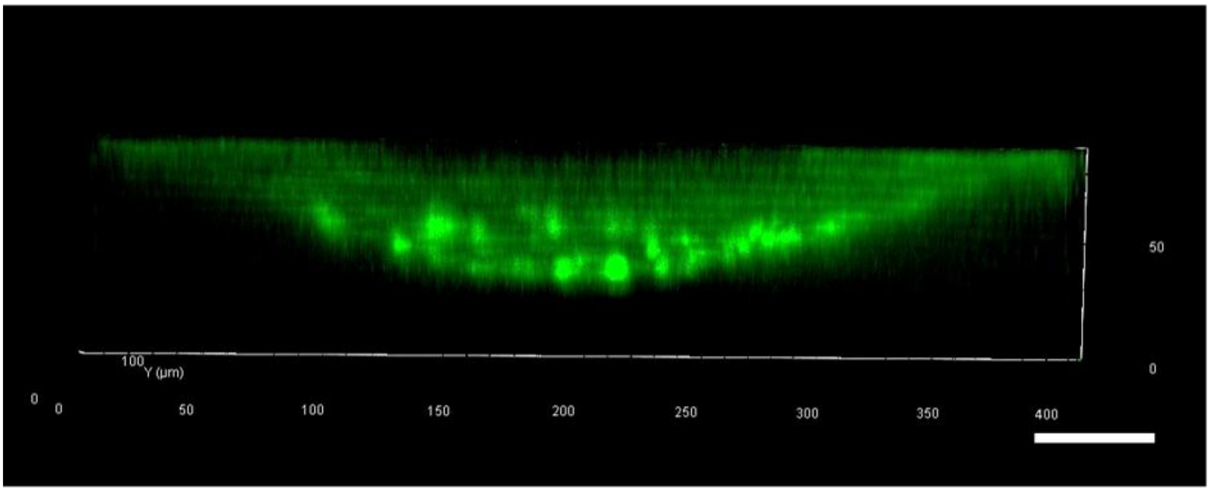


**Supplementary Figure 6.** Side-view fluorescence image of FITC-labeled alveoli-mimetic PLGA membranes. Scale bar: 50 μm.

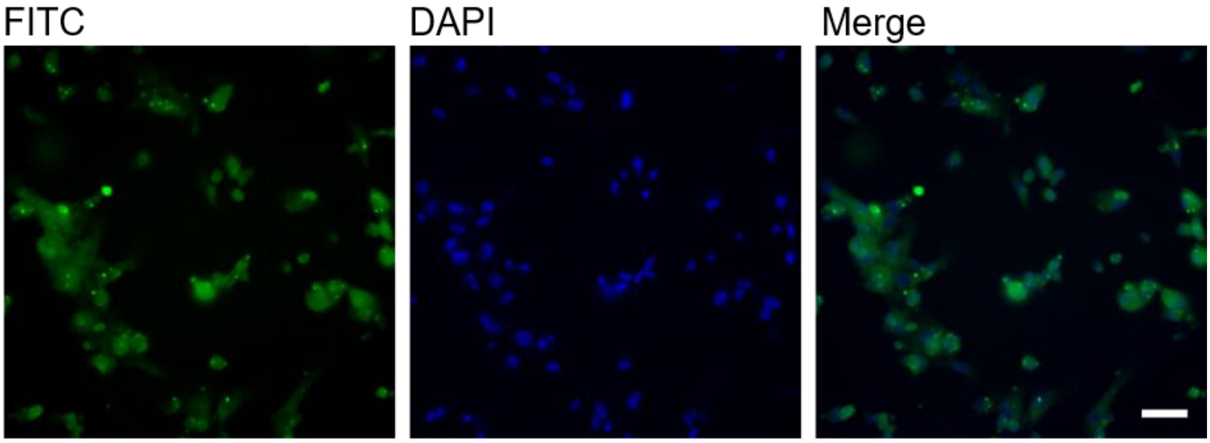


**Supplementary Figure 7. Cellular uptake of FITC-labeled PLGA by lung fibroblasts.** Immunofluorescence images showing lung fibroblasts cultured on FITC-labeled PLGA membranes for 5 days. Cells were trypsinized prior to imaging to assess intracellular uptake of PLGA. Scale bar: 50 μm.

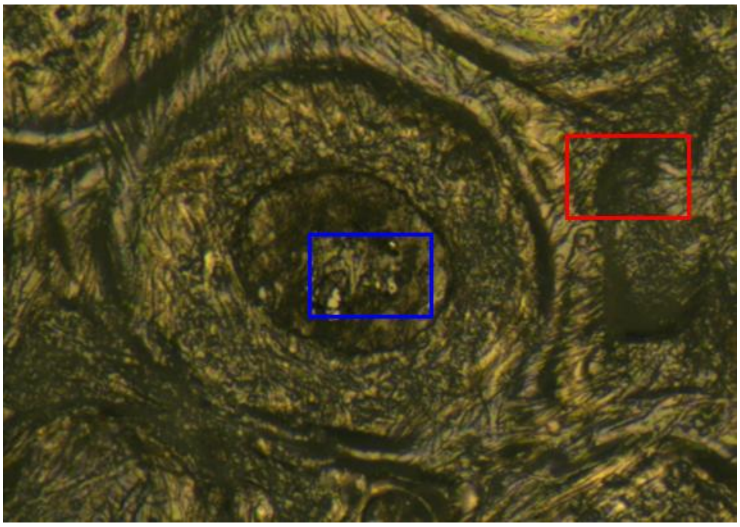

**Supplementary Figure 8.** Bright-field image showing fibroblasts cultured on a PLGA membrane after degradation. The blue boxed region indicates areas where the PLGA membrane has degraded, leaving only the cell-derived layer, whereas the red boxed region indicates areas with remaining PLGA membrane. Raman spectra were acquired from both the blue- and red-boxed regions.

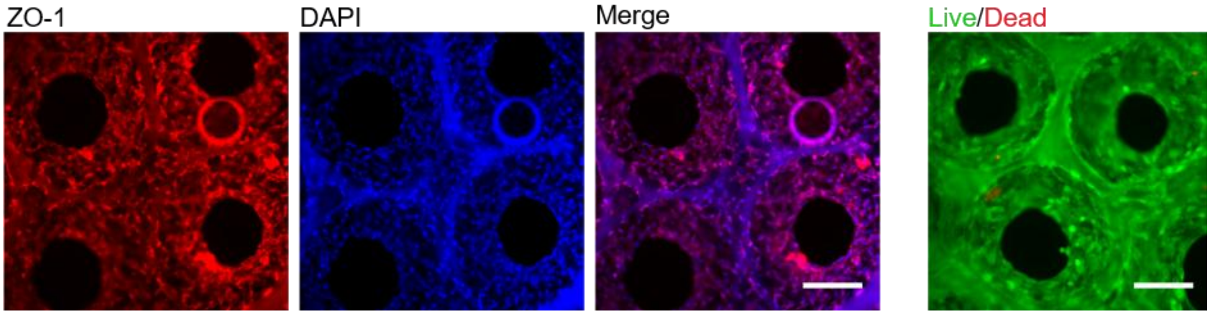


**Supplementary Figure 9**. **A549 monoculture on the chip in the absence of lung fibroblasts.** A549 cells were cultured on the chip for 7 days without lung fibroblasts. The left panel shows immunofluorescence staining of ZO-1 (red), with nuclei counterstained with DAPI (blue). The right panel shows live/dead staining images. Scale bars: 200 μm.

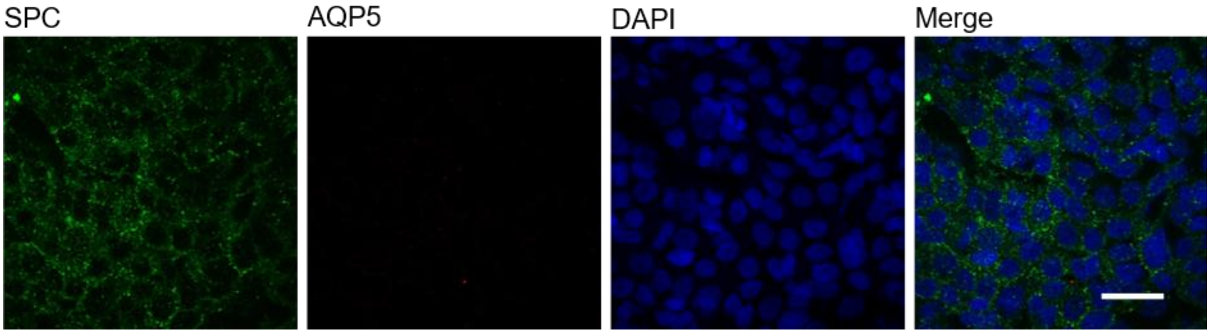


**Supplementary Figure 10**. **Alveolar epithelial phenotype validation.** A549 cells cultured on the chip were stained for aquaporin 5 (AQP5, red), surfactant protein C (SPC, green), and nuclei (DAPI, blue). Scale bar: 50 µm.

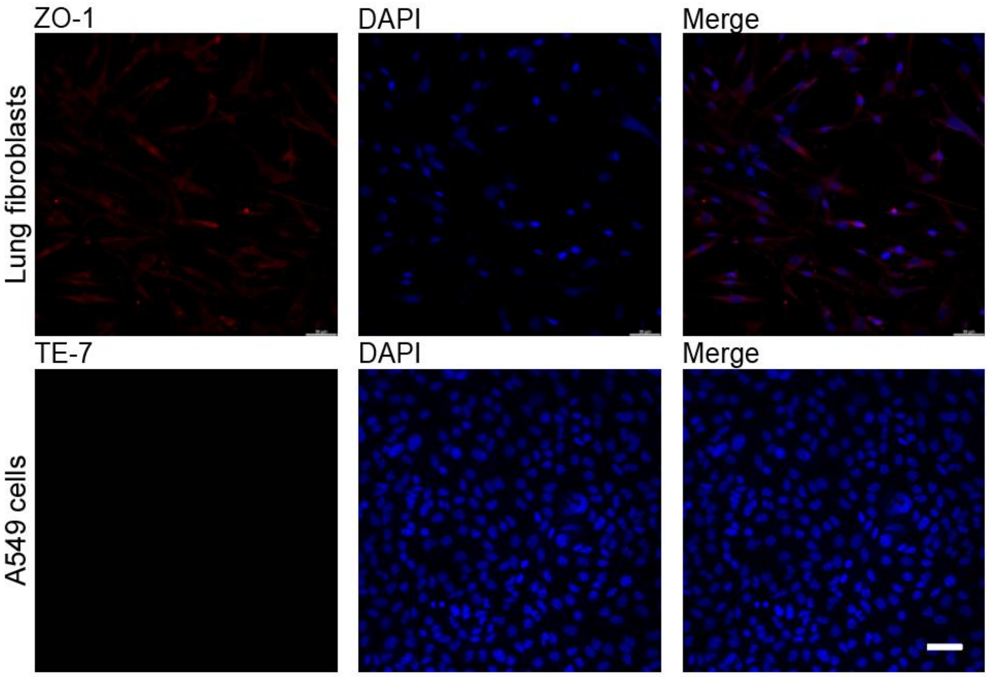


**Supplementary Figure 11. Validation of cell-type-specific markers in lung fibroblasts and A549 cells.** Immunofluorescence staining of lung fibroblasts for ZO-1 (top) and A549 cells for TE-7 (bottom). No detectable signal was observed in either case, confirming the specificity of epithelial and fibroblast markers. Nuclei were counterstained with DAPI (blue). Scale bar: 50 µm.

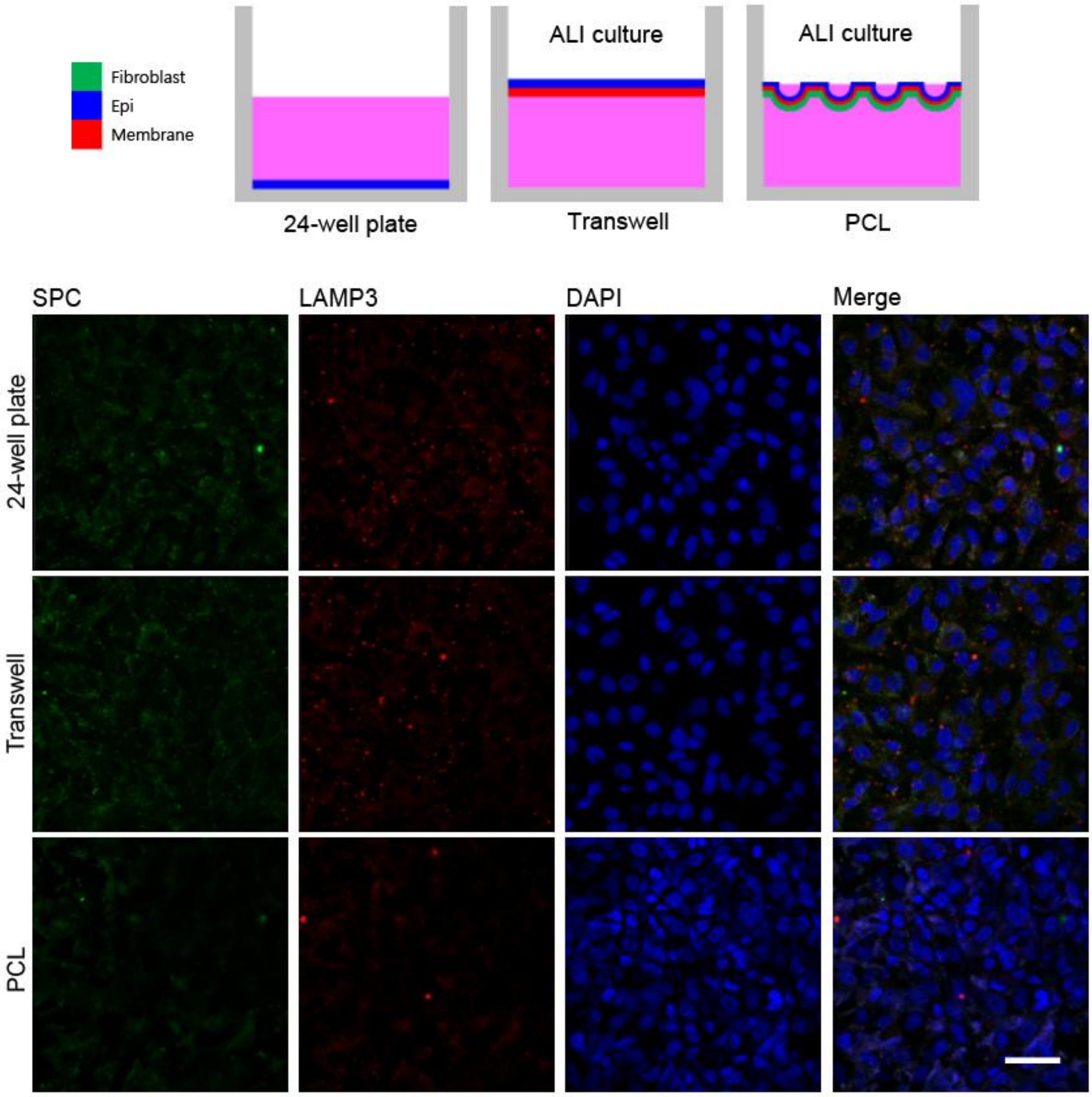


**Supplementary Figure 12**. Immunofluorescence images of A549 cells cultured on a 24-well plate, Transwell®, and PCL membrane, stained for SPC, LAMP3, and DAPI. ALI culture was applied in the Transwell® and PCL membrane conditions. Scale bar: 50 μm.

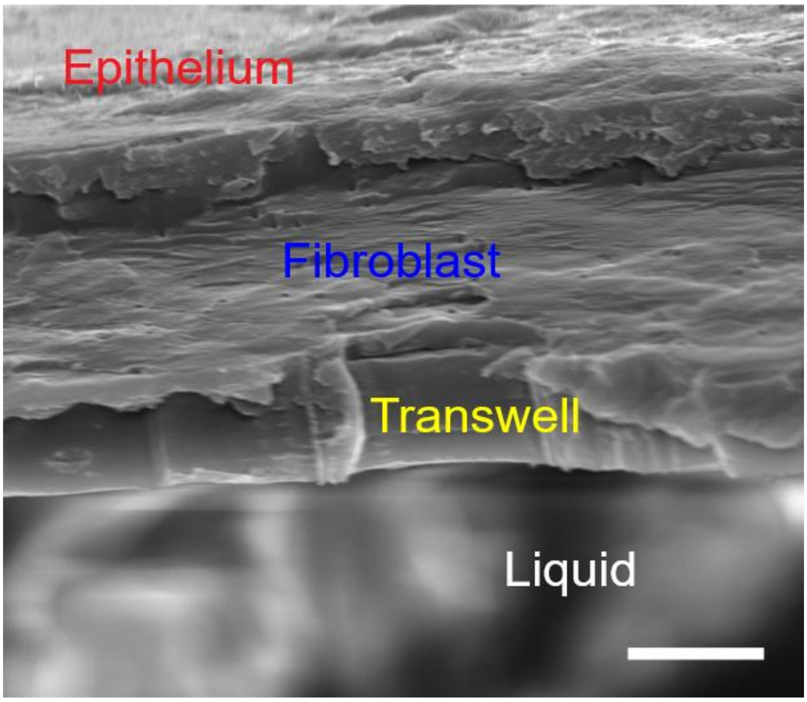


**Supplementary Figure 13**. Cross-sectional SEM image of epithelial–fibroblast co-culture on a Transwell® membrane. Lung fibroblasts were cultured on a Transwell® membrane for 2 days, followed by seeding of A549 cells on top. A549 cells were maintained for 5 days, including 2 days under ALI conditions. Scale bar: 10 µm.

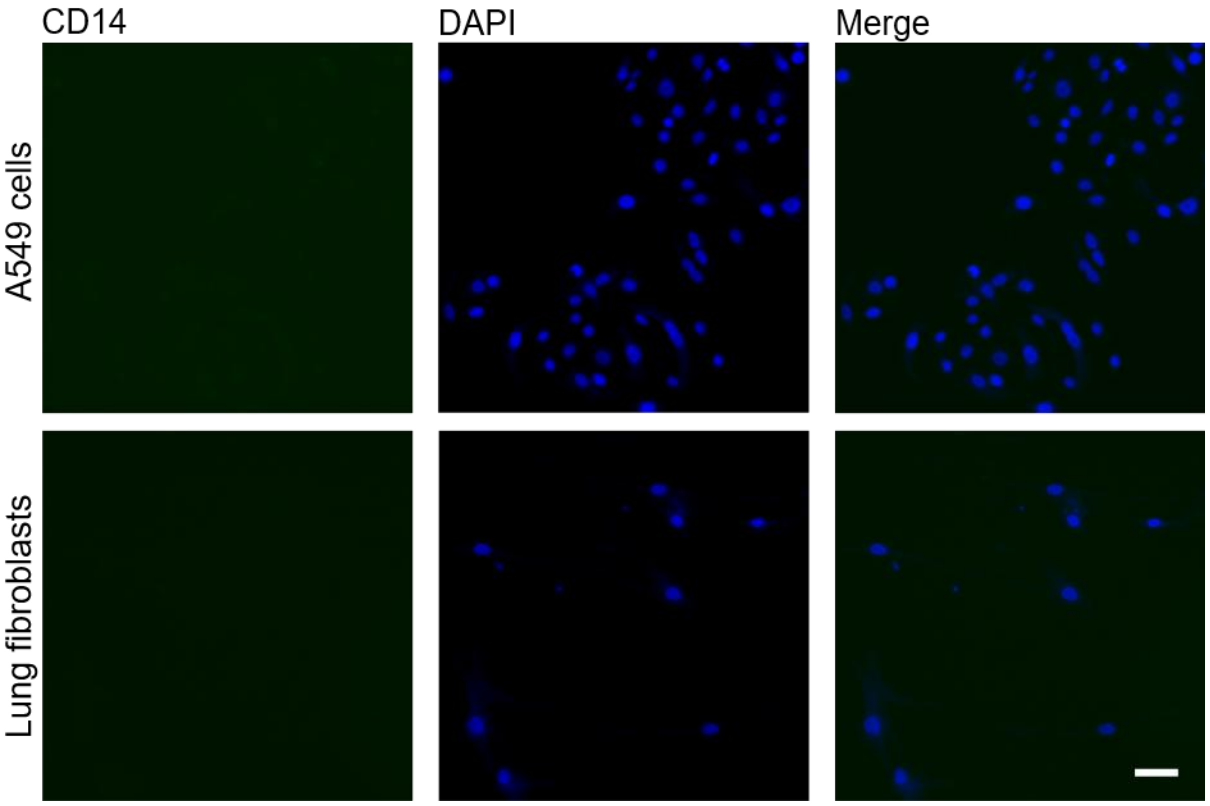


**Supplementary Figure 14**. Immunofluorescence images of A549 cells and lung fibroblasts stained for CD14 (green), with nuclei counterstained with DAPI (blue). No detectable CD14 expression was observed in either cell type. Scale bar: 50 µm.

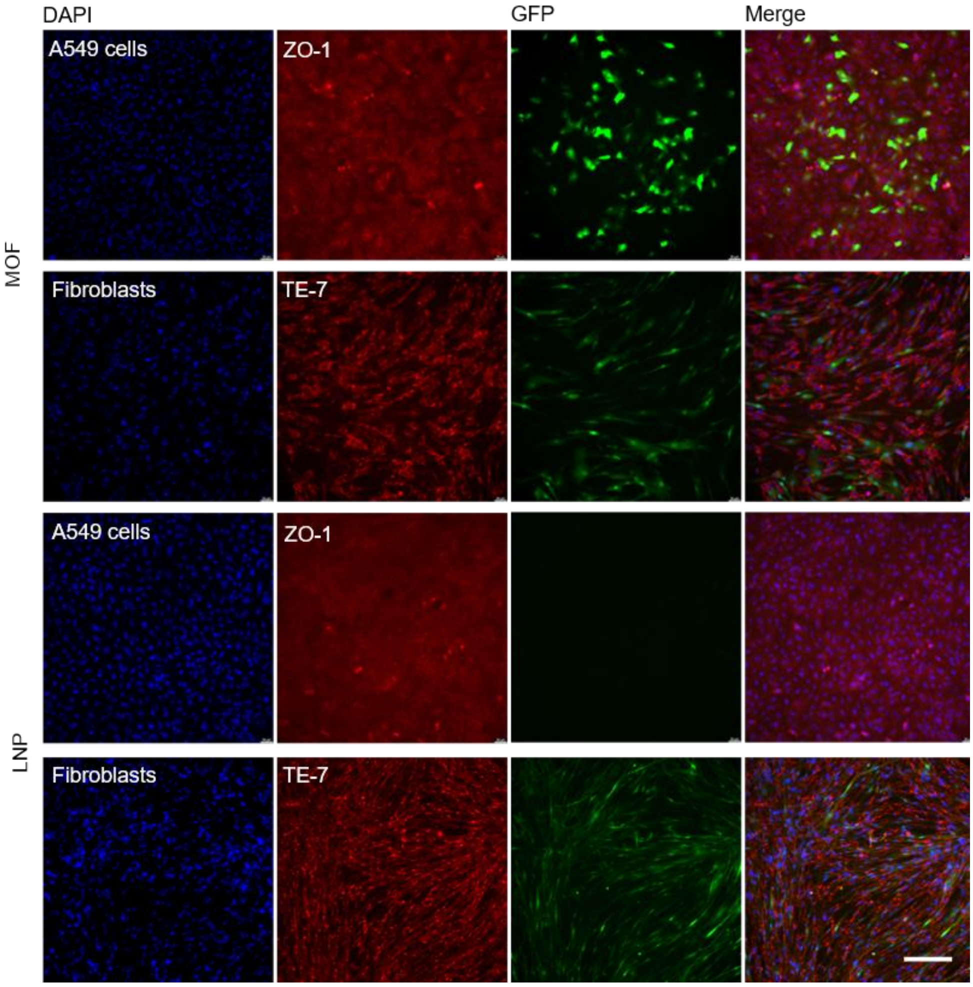


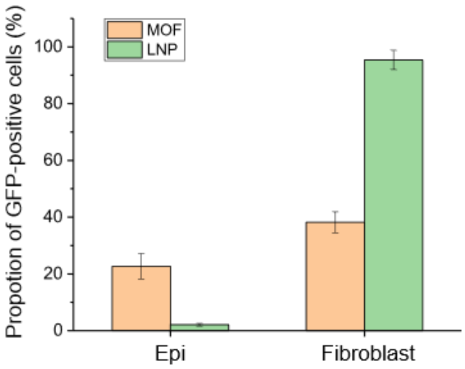


**Supplementary Figure 15**. **Comparison of MOF and LNP transfection efficiency in A549 cells and lung fibroblasts.** (Top) Immunofluorescence images of A549 cells and lung fibroblasts cultured separately in 6-well plates and exposed to MOF or LNP for 48 h. A549 cells were stained with ZO-1 and fibroblasts with TE-7, with nuclei counterstained by DAPI (blue). Green fluorescence indicates successful transfection. Scale bar: 200 μm. (Bottom) Quantification of transfection efficiency in A549 cells and lung fibroblasts following 48 h exposure to MOF or LNP (n = 4).

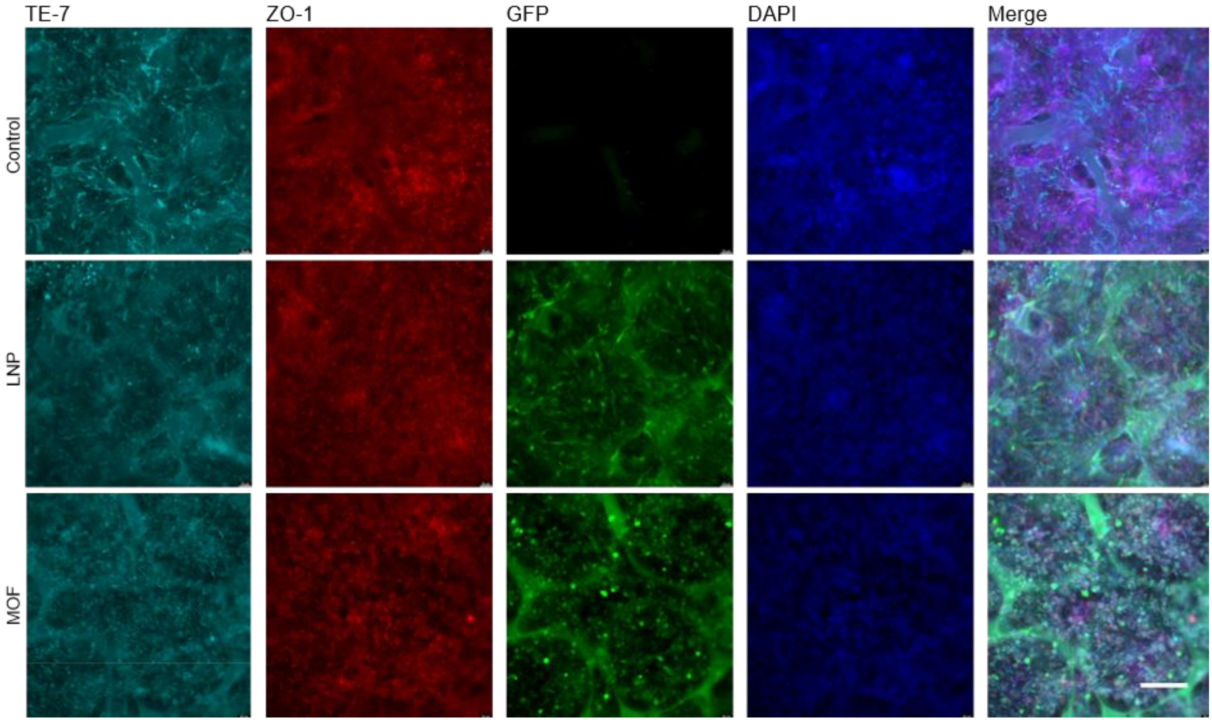


**Supplementary Figure 16.** Immunofluorescence images showing cellular responses in the alveoli-on-a-chip following nanoparticle delivery in the absence of macrophages. Cells were stained for fibroblasts (TE-7), epithelial tight junctions of A549 cells (ZO-1), GFP expression indicating nanoparticle-mediated gene delivery, and nuclei (DAPI). Images correspond to control, LNP-treated (1 µg, 48 h), and MOF-treated (1 µg, 48 h) conditions. Scale bar: 200 µm.

| **Antibody** | **Company** | **Cat. No.** | **Dilution** |
|---|---|---|---|
| Anti-Fibroblasts Antibody, clone TE-7 | Sigma-Aldrich | CBL271 | 1:200 |
| Anti-ZO-1 Antibody | Rockland | 600-401-GU7 | 1:400 |
| Fibronectin Polyclonal antibody | Proteintech | 15613-1-AP | 1:200 |
| LAMP3 Polyclonal antibody | | 12632-1-AP | 1:400 |
| Alpha smooth muscle actin specific Polyclonal antibody | | 55135-1-AP | 1:500 |
| COL1A Antibody | Santa Cruz | sc-59772 | 1:200 |
| Surfactant protein c Antibody | | sc-518029 | 1:200 |
| Aquaporin 5 Antibody | | sc-514022 | 1:200 |
| CD14 Monoclonal Antibody | Invitrogen | 14-0149-82 | 1:200 |
| CD68 Monoclonal Antibody | | 14-0688-82 | 1:200 |
| Phalloidin-iFluor 647 Reagent | Abcam | ab176759 | 1:1000 |
| Surfactant protein c Antibody | Seven Hills Bioreagents | WRAB-9337 | 1:200 |

**Table S1.** List of antibodies.
